# TRMT1L-catalyzed m^2^_2_G27 on tyrosine tRNA is required for efficient mRNA translation and cell survival under oxidative stress

**DOI:** 10.1101/2024.05.02.591343

**Authors:** Sseu-Pei Hwang, Han Liao, Katherine Barondeau, Xinyi Han, Cassandra Herbert, Hunter McConie, Amirtha Shekar, Dimitri Pestov, Patrick A Limbach, Jeffrey T Chang, Catherine Denicourt

**Affiliations:** Department of Integrative Biology and Pharmacology, McGovern Medical School, The University of Texas Health Science Center, Houston, TX 77030, USA; The University of Texas MD Anderson Cancer Center UTHealth Houston Graduate School of Biomedical Sciences, Houston, TX 77030, USA; Rieveschl Laboratories for Mass Spectrometry, Department of Chemistry, University of Cincinnati, Cincinnati, OH 45221, USA; Department of Cell Biology and Neuroscience, Virtua Health College of Medicine and Life Sciences, Rowan University, Stratford, NJ 08028, USA

**Keywords:** TRMT1L, RNA methyltransferases, N2, N2-dimethylguanosine (m^2^_2_G), eCLIP-seq, Nanopore tRNA-seq, tRNA, rRNA, translation, oxidative stress response

## Abstract

tRNA modifications are critical for several aspects of their functions, including decoding, folding, and stability. Using a multifaceted approach encompassing eCLIP-seq and Nanopore tRNA-seq, we show that the human tRNA methyltransferase TRMT1L interacts with component of the Rix1 ribosome biogenesis complex and binds to the 28S rRNA, as well as to a subset of tRNAs. Mechanistically, we demonstrate that TRMT1L is responsible for catalyzing m^2^_2_G solely at position 27 of tRNA-Tyr-GUA. Surprisingly, TRMT1L depletion also impaired the deposition of acp^3^U and dihydrouridine on tRNA-Tyr-GUA, Cys-GCA, and Ala-CGC. TRMT1L knockout cells have a marked decrease in tRNA-Tyr-GUA levels, coinciding with a reduction in global translation rates and hypersensitivity to oxidative stress. Our results establish TRMT1L as the elusive methyltransferase catalyzing the m^2^_2_G27 modification on tRNA Tyr, resolving a long-standing gap of knowledge and highlighting its potential role in a tRNA modification circuit crucial for translation regulation and stress response.

## INTRODUCTION

Post-transcriptional modification of RNA molecules has emerged as a key regulatory component of eukaryotic gene expression.^1,2^ Around 170 different types of chemical modification have been reported on coding and non-coding RNAs from all three domains of cellular life.^2^ In eukaryotes, nuclear-encoded tRNAs are the most extensively modified RNA, containing on average 10-13 modifications per molecule.^3^ tRNAs are essential adapter molecules that deliver amino acids to the ribosome to translate the genetic information encoded by mRNA templates.^4^ Modification expands the chemical properties of RNA nucleosides, thereby having a direct influence on tRNA structure, stability, and decoding function.^3,5^ Proper modification of tRNAs is essential for accurate and efficient translation and as such, the deregulation of tRNA modification and the enzymes responsible for their catalysis have been linked with altered mRNA translation programs in several human diseases.^6,7^

TRMT1L is a poorly characterized vertebrate paralog of the evolutionarily conserved tRNA methyltransferase, TRMT1.^8,9^ TRMT1 catalyzes the deposition of N2, N2-dimethylguanosine (m^2^_2_G) at position 26 of cytosolic and mitochondrial tRNAs.^9–14^ Unlike TRMT1, which possesses a mitochondrial-targeting sequence and localizes to mitochondria, cytosol, and nucleus, TRMT1L predominantly resides in the nucleus and nucleolus.^8,9^ Noteworthy is the shared significance of TRMT1 and TRMT1L in regulating neuronal functions, as studies demonstrate that genetic deletion of TRMT1L in mice disrupts motor coordination and induces aberrant exploratory behaviors.^8,9,15–17^ While the activity and targets of TRMT1 have been extensively characterized across various organisms, including humans, the substrates and function of TRMT1L remain incompletely elucidated. Recent investigations examining mouse brains RNA-seq datasets for modifications-induced mismatch signatures, suggested that TRMT1L could catalyze the deposition of m^2^_2_G at position 26 on tRNA-Ala.^8^ These findings imply that, akin to TRMT1, TRMT1L may function as an m^2^_2_G RNA methyltransferase, potentially evolving to act upon a similar or distinct set of substrates.

The m^2^_2_G-modified nucleotide at position 26 is found in the structural hinge region formed between the dihydro-uridine stem (D-stem) and the anticodon stem loop (ASL) in the vast majority of eukaryotic tRNAs.^18,19^ The presence of a N2, N2 dimethyl group on G precludes its pairing with C and, consequently m^2^_2_G pairs almost exclusively with A, U, or G at position 44 of tRNAs. Consistent with the crucial roles of modification in regulating tRNA structure, both molecular dynamic simulations and crystal structure determination studies revealed that m^2^_2_G not only inhibits the pairing of G26 with C in the tRNA D-stem but also regulates the pairing mode with A, favoring a more stable imino-hydrogen bonded form.^20–22^ The m^2^_2_G modification undergoes dynamic regulation in response to diverse stress.^23,24^ Studies in human cells have revealed that depletion of TRMT1 results in diminished global mRNA translation and increased susceptibility to oxidizing agents.^9^ These findings suggest a crucial role for m^2^_2_G modification in maintaining protein synthesis efficiency and cellular redox balance.

Here, we report the functional characterization of TRMT1L in human cells. While performing a proteomic purification of the Rix1 60S ribosome biogenesis complex, we identified TRMT1L as an interacting component. Using an eCLIP-seq approach, we demonstrate that TRMT1L specifically binds to the 28S rRNA expansion segment ES7Lb and to a small subset of nuclear-encoded tRNAs, of which Cys-GCA and Tyr-GUA were the most abundantly enriched. Using CRISPR-generated TRMT1L knock-out (KO) cells, we find that TRMT1L is required to catalyze the deposition of m^2^_2_G at position 27 of only tRNAs Tyr-GUA. TRMT1L-dependent m^2^_2_G modifications were not detected of any other tRNAs nor the 28S rRNA, suggesting that it may have catalytically independent functions depending on the RNA substrate. Intriguingly, TRMT1L-deficiency also leads to a loss of 3-(3-amino-3-carboxypropyl) uridine (acp^3^U) and dihydrouridine (D) modifications on Tyr-GUA, Cys-GCA and Ala-CGC tRNAs, suggesting that TRMT1L may be part of a modification circuit required for the deposition of acp^3^U and D by DTWD1 and DUS enzymes, respectively. Quantitative analysis of tRNA abundance in TRMT1L KO cells revealed a significant decrease in tRNA-Tyr-GUA levels, which correlated with a decline in global mRNA translation and hypersensitivity to endoplasmic reticulum (ER) and oxidative stress compared to WT cells. Our findings reinforce the importance of m^2^_2_G in tRNA regulation for stress response and suggest an essential role for TRMT1L in regulating tRNA functions and establishing an adaptive translational program in response to oxidative and ER stress.

## RESULTS

### TRMT1L interacts with components of the Rix1 complex and co-sediments with pre-60S particles

In a mass spectrometry-based proteomic analysis of the Rix1 60S biogenesis complex through immunoprecipitation of endogenous PELP1, we identified the RNA methyltransferase TRMT1L as an interacting component (Figure 1A). Immunoprecipitation of PELP1 followed by TRMT1L Western blot analysis was used to validate this interaction (Figure 1B). Reversely, we also show that immunoprecipitated endogenous TRMT1L interacts with PELP1 and other components of the larger Rix1 complex, namely WDR18 and SENP3 and with the ribosome biogenesis factor NPM1 (Figure 1C).^25,26^ TRMT1L has previously been reported to localize to the nucleus, with an enrichment to the nucleolar compartment, which is the site of ribosome biogenesis.^8,9^ Immunostaining analysis on U2OS cells treated with control and TRMT1L siRNAs validated the localization of endogenous TRMT1L in the nucleolus (Figure 1E). We predicted putative nucleolar (NoLS) and nuclear (NLS) localization sequences situated at the N- and C-terminus of TRMT1L, respectively (Figure 1D and S1A).^27–29^ Mutation of key basic residues within the NoLS domain resulted in a redistribution of TRMT1L in the nucleoplasm. In contrast, mutation of basic residues in the NLS domain resulted in cytosolic redistribution of TRMT1L, indicating these regions act as critical regulatory elements governing its subcellular distribution (Figure S1B-C). Interestingly, the nucleolar localization of TRMT1L appears to depend on active rRNA transcription as inhibition of RNA Pol I activity with CX-5461 resulted in its redistribution to the nucleoplasm (Figure 1F).

**Figure 1.**
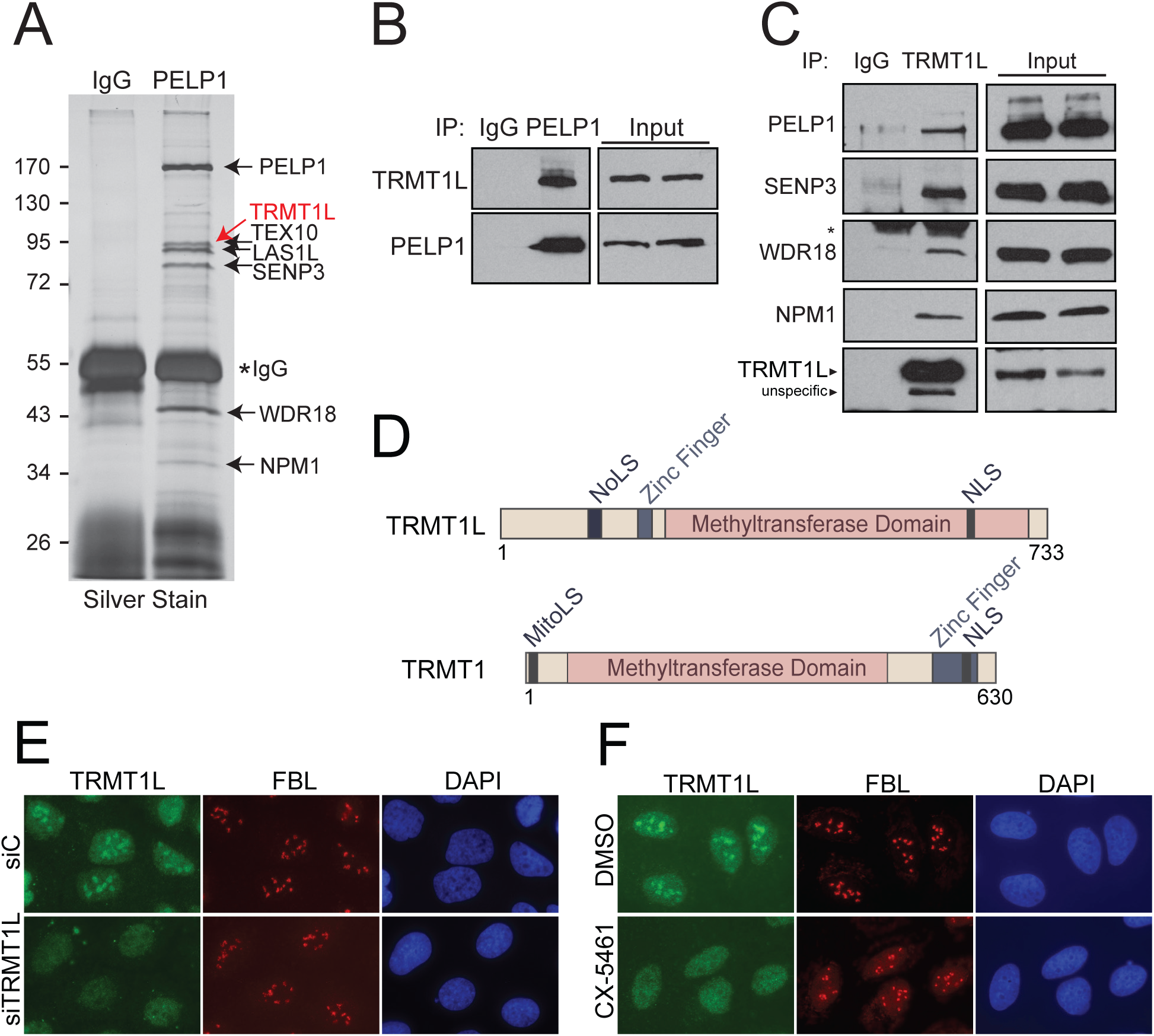
TRMT1L interacts with components of the Rix1 60S biogenesis complex in nucleolus. **(A)** Mass spectrometry analysis of PELP1-associated complexes. **(B)** Western blot analysis of endogenous PELP1 or **(C)** TRMT1L immunoprecipitated (IP). Asterisk indicates the presence of the IgG cross-reactivity band. **(D)** Schematic representation of TRMT1L and TRMT1 proteins. **(E)** Immunofluorescence detection of endogenous TRMT1L. **(F)** Inhibition of rRNA transcription by CX-5461 impairs TRMT1L nucleolar localization in U2OS cells. See also Figure S1.

PELP1 and the Rix1 complex regulate important rRNA processing steps required for 60S ribosomal subunit biogenesis.^25,26^ As we co-purified TRMT1L with Rix1 components, we tested whether TRMT1L regulates ribosome synthesis. We first determined whether TRMT1L co-sediments with nucleolar pre-ribosomal particles by sucrose gradient sedimentation profiling. Similarly to PELP1 and WDR18, we find that TRMT1L specifically co-sediment with only the pre-60S subunit (Figure 2A). However, upon TRMT1L depletion by RNAi, we did not observe any significant rRNA processing defects apart from a slight reduction in the levels of the 32S and 12S rRNA precursors (Figure S2A-C). Nuclear 40S and 60S pre-ribosomal particle levels were also not affected by TRMT1L depletion (Figure 2B-C). Collectively, these findings suggest that while TRMT1L co-sediments with pre-60S ribosomes, it is unlikely to be directly involved in rRNA processing or be indispensable for ribosome biogenesis. This observation is consistent with the limited evolutionary conservation of TRMT1L and the viability of TRMT1L gene knockout mice,^17^ suggesting a regulatory rather than essential role for TRMT1L in ribosome function.

**Figure 2.**
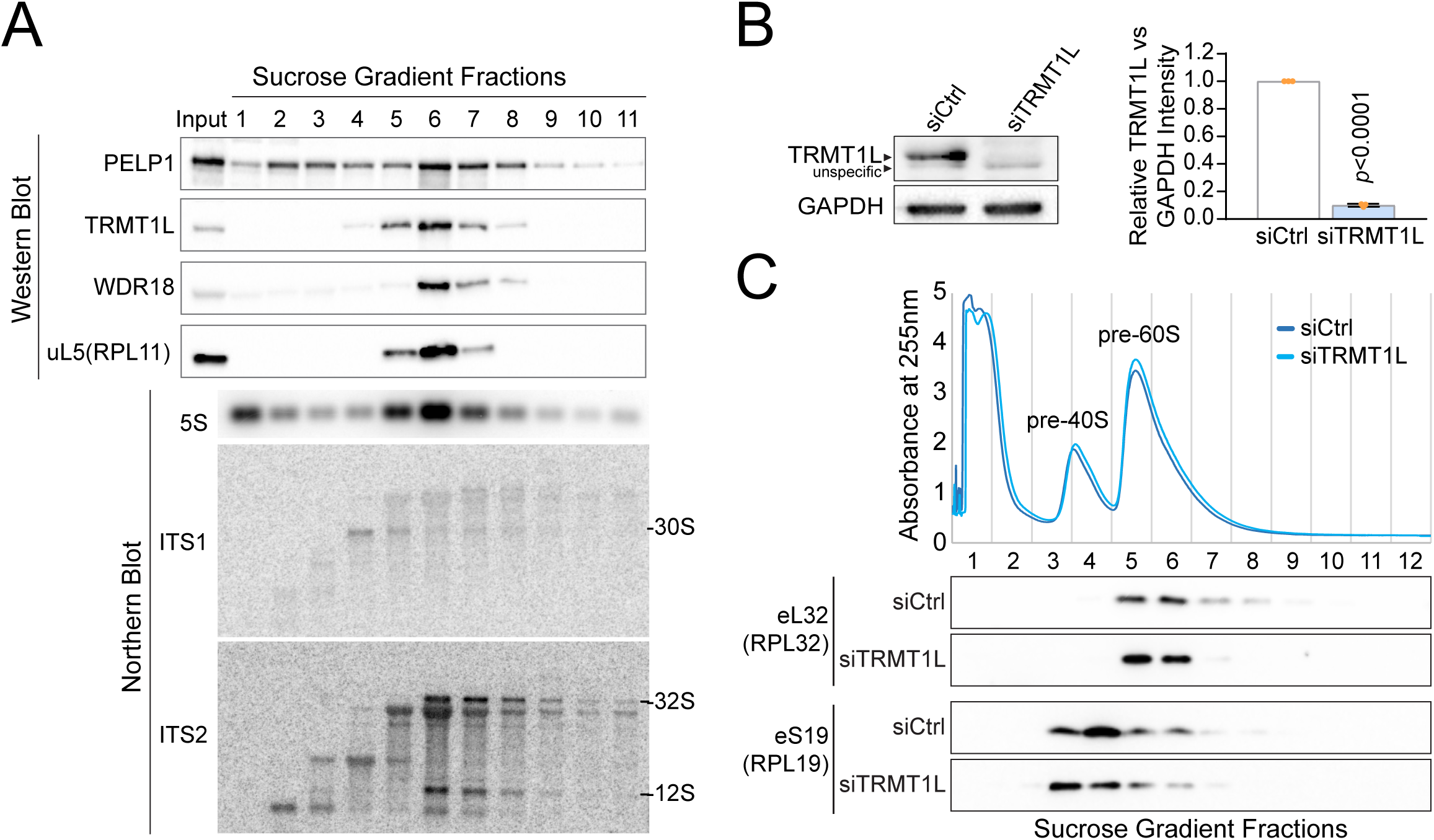
TRMT1L co-sediments with pre-60S ribosomes. **(A)** Pre-ribosomal particles from sucrose gradient fractionation. Proteins and RNA from each fraction were analyzed by Western blot (top panel) or by Northern blot (bottom panel), respectively. **(B)** RNAi depletion of TRMT1L was confirmed by immunoblotting and densitometry quantification Data are presented as mean ± SD. N=3 **(C)** Nuclear pre-ribosomes from TRMT1L depleted cells were separated by sucrose gradient fractionation and analyzed by Western blot. See also Figure S2.

### TRMT1L binds to tRNAs and 28S rRNA expansion segments

To further elucidate TRMT1L functions and identify its potential RNA substrates, we performed a genome-wide eCLIP-seq analysis^30,31^ of endogenous TRMT1L in two different cell lines (HCT116 and HeLa) in duplicates and with respective size-matched input (SMInput) controls. We first tested whether TRMT1L binds to RNA by performing a ligation-mediated labeling of immunoprecipitated (IP) endogenous TRMT1L with pCp-Biotin.^32^ The pCp-Biotin signal was detected only in the cross-linked TRMT1L IP, not in the non-crosslinked condition or the control IgG IP, indicating the presence of trapped TRMT1L-RNA complexes (Figure S3A-B). Having validated the binding of TRMT1L to RNA, cells were processed for large-scale eCLIP-seq analysis, as depicted in Figure S3C.

Most eCLIP reads in TRMT1L samples corresponded to rRNAs and tRNAs (Figure 3A and Table S1), which also showed peaks of significant enrichments when normalized to SMInput (Figure 3B-C, Figure S4). For tRNAs, the majority of enriched reads in both HCT116 and HeLa cells corresponded to a small subset of nuclear-encoded tRNAs, mainly tRNA-Cys-GCA, Tyr-GUA followed by Arg-UCG, Arg-CCU, and Asn-GUU genes (Figure 3B-C). Although in lower abundance, other various isoacceptors were found in HCT116 vs. HeLa cells differently, indicating that the range of tRNA substrates is cell-specific (Figure 3B-C and Figure S4A-B). Remarkably, the reads enriched in the TRMT1L immunoprecipitation (IP) for tRNA-Cys-GCA and Tyr-GUA corresponded to precursor tRNAs that retained their 3’ trailer sequence. In contrast, most other tRNAs appeared mature, with their 3’ trailer processed and containing the CCA addition (Figure S4C). For tRNA-Tyr-GUA, most reads retaining the 3’ trailer exhibited intron removal, suggesting that TRMT1L binds to these tRNAs during an early processing stage. Surprisingly, for rRNA, a main peak of enrichment aligned to a very narrow region of the 28S rRNA corresponding to expansion segments ES7Lb (Figure S4D). In conclusion, our CLIP-seq analysis indicates that TRMT1L binds to a specific subset of tRNAs and the 28S rRNA expansion segment ES7Lb.

**Figure 3.**
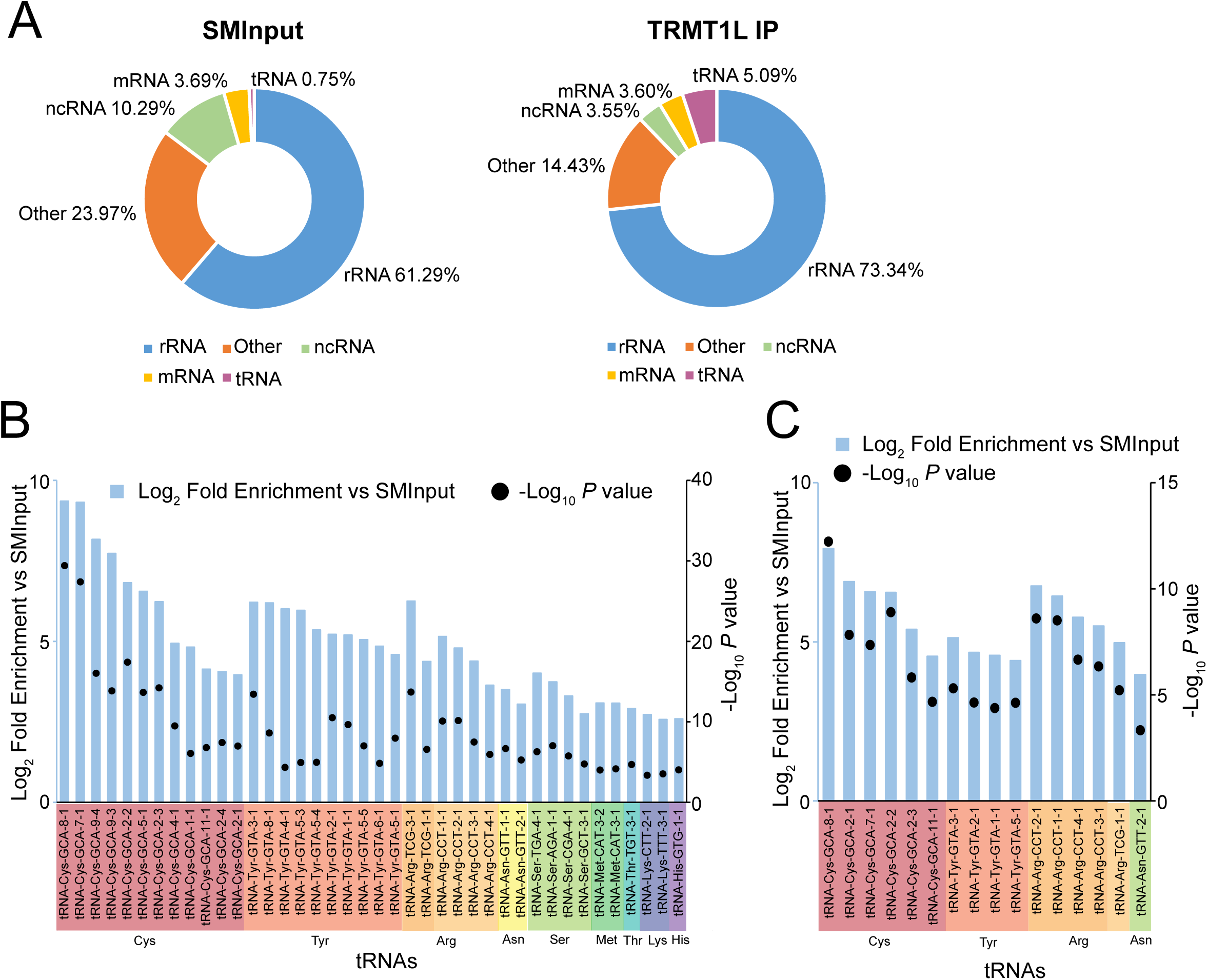
TRMT1L binds to tRNAs and rRNA. **(A)** eCLIP-sequencing analysis showing percentage of reads in SMInputs and TRMT1L IP samples mapped to CLIP peak regions for HCT116 cells. **(B)** Log_2_ fold enrichment of TRMT1L IP over SMInput with adjusted p values < 0.05, fold change ≥ 2, and basemean ≥ 1000 from HCT116 or **(C)** HeLa duplicates for different tRNA iso-decoders. See also Figure S3 and S4 and Table S1.

### TRMT1L catalyzes the deposition of m^2^_2_G at position 27 of tRNA-Tyr-GUA

To test the presence of TRMT1L-catalyzed modification on rRNA, we established CRISPR Cas9-mediated TRMT1L knockout HCT116 cells with two different guide RNAs (TRMT1L KO1 and KO2) (Figure S5A-C). As expected, immunoblotting revealed the absence of detectable TRMT1L protein in both KO1 and KO2 cell lines. Importantly, we did not observe any significant effect on the levels of the paralog TRMT1, indicating the suitability of these cell lines to specifically assess TRMT1L functions (Figure S5C). We next isolated 28S rRNA from WT (non-targeting control guide RNA) and TRMT1L KO1 cells (Figure S5D) for quantitative liquid chromatography-tandem mass spectrometry (LC-MS/MS) of modified ribonucleosides. Among the reported 28S modification levels measured, in addition to m^2^G and m^2^_2_G, no significant differences were detected in the TRMT1L KO compared to the control WT RNA samples (Figure S5E). Importantly, m^2^G and m^2^_2_G modifications were not detected on 28S rRNA. This observation aligns with previous LC-MS/MS analysis,^33^ which did not uncover any m^2^G or m^2^_2_G on human ribosomes.

To evaluate the occurrence of TRMT1L-catalyzed modifications on tRNAs, a significant potential substrate revealed in the eCLIP-seq analysis, we initially employed a Northern blot-based PHA approach (Positive Hybridization in Absence of modification),^34,35^ comparing total RNA from WT and TRMT1L KO cells. This assay relies on the differential hybridization efficiency of a probe due to the presence or absence of a bulky modification, which affects base pairing. We detected an elevated PHA signal exclusively for tRNAs Cys-GCA and Tyr-GUA from the TRMT1L KO cells (Figure S5F-J). This signal was observed when using a probe covering the D-loop side of the tRNAs, while no such signal was detected with a probe pairing with the T-loop side, which served as a normalization control (Figure S5H and J). Other tRNA tested did not show any PHA effect in the TRMT1L KO cells (Figure S5I-J). This suggests a potential loss of modification on the D-loop side of tRNAs Cys-GCA and Tyr-GUA. Notably, ectopic re-expression of TRMT1L in the KO background markedly rescued the heightened PHA signal on both tRNA-Cys-GCA and Tyr-GUA, implying that the potential loss of modification is TRMT1L-dependent (Figure S5K-M). However, a catalytically inactive TRMT1L mutant (D373V), which localizes to the nucleolus similarly to WT (Figure S6A), did not rescue the increased PHA signal on tRNA Tyr-GUA. Interestingly, on tRNA-Cys-GCA, re-expression of the D373V mutant was able to rescue the PHA signal to the same extent as TRMT1L WT, indicating that TRMT1L may have catalytically independent activities, perhaps assisting another RNA modifying enzyme to perform its function (Figure S5K-M). Our results indicate that TRMT1L likely regulates modification on the D-loop side of tRNAs Cys-GCA and Tyr-GUA.

To uncover TRMT1L-catalyzed modifications on tRNAs, we employed Nano-tRNAseq, a Nanopore-based direct RNA sequencing method.^36^ Direct RNA nanopore sequencing detects RNA modifications with near-single nucleotide precision by identifying positions showing base calling errors upon alignment to a reference transcriptome.^36–45^ Purified tRNAs isolated from WT and TRMT1L KO cells were processed for Nano-tRNAseq and the base calling errors (base mismatch, insertion and deletion) for each nucleotide position and tRNA iso-decoder reference were calculated for each duplicate sample. Relative to WT, TRMT1L KO cells showed a significant decrease in base calling errors for specific positions mainly for tRNA-Tyr-GUA and Cys-GCA (Figure 4A), the top two iso-decoders found to have the highest enrichment for binding to TRMT1L in the eCLIP-Seq analysis (Figure 3B-C). Although not found to be significantly enriched in the TRMT1L eCLIP-Seq data, Ala-CGC and Ala-UCG also showed positions with decreased base calling errors (Figure 4A), suggesting the loss of modification on these tRNAs in the TRMT1L KO cells. A closer examination of specific nucleotide positions with decreased base calling errors in the TRMT1L KO revealed potential loss of m^2^_2_G27 and surprisingly 3-(3-amino-3-carboxypropyl) uridine (acp^3^U) 20 and dihydrouridine (D) 17, modifications reported to be present at these positions on tRNA-Tyr-GUA.^46,47^ Both tRNAs Cys-GCA and Ala-CGC/UCG from TRMT1L KO cells showed a significant increase in quality reads at position 20, suggesting loss of acp^3^U that were reported on these tRNAs.^46,48^ In addition, we also observed a slight increase in quality reads at position 16 for tRNA-Ala-CGC/UCG, which could correspond to the loss of D at this position (Figure 4B-C).

**Figure 4.**
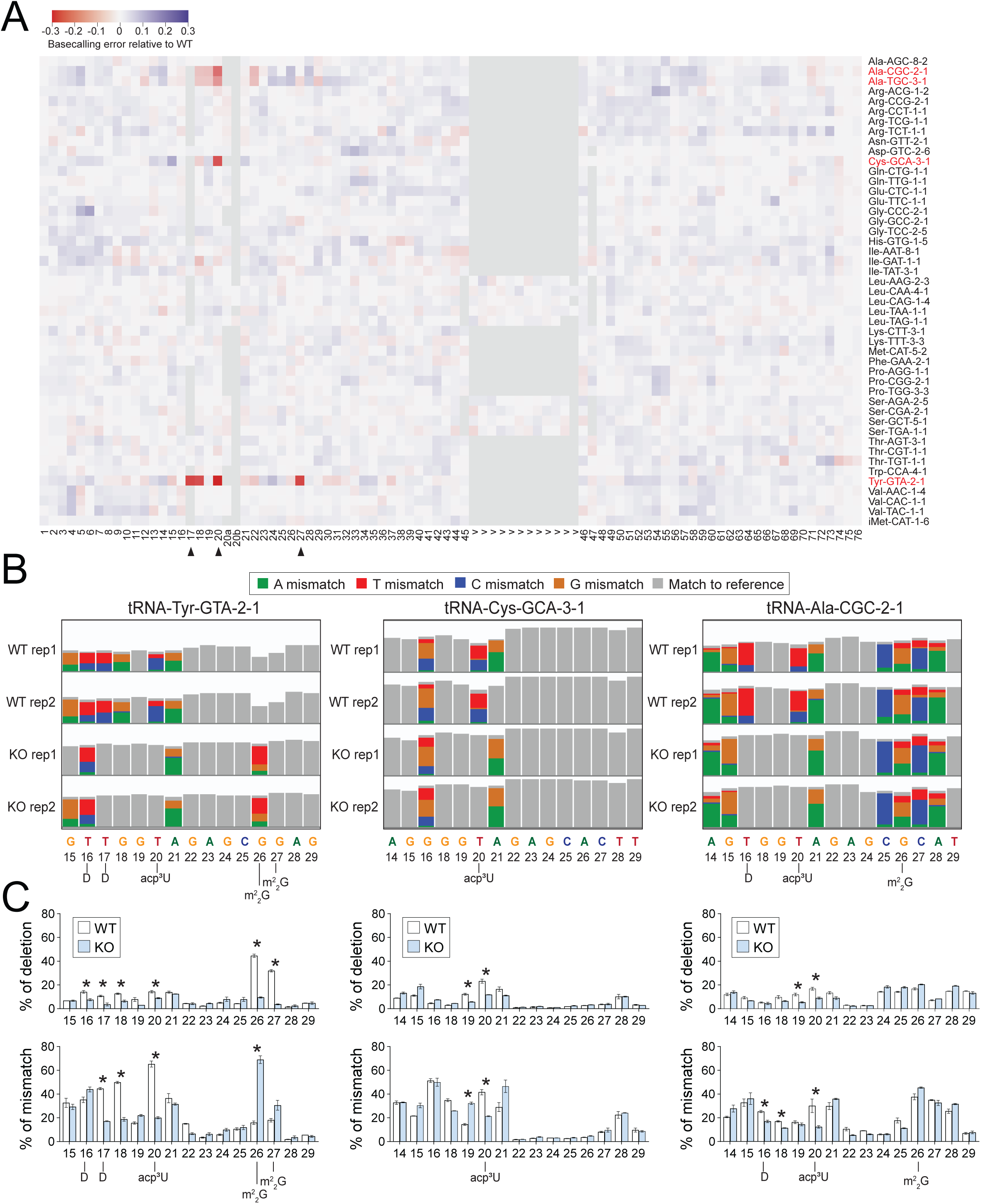
Nano-tRNAseq identifies potential tRNA modification changes in TRMT1L-depleted cells. **(A)** Heatmap of summed base calling errors in TRMT1L KO cells vs. WT, for each nucleotide and tRNA. Triangles mark positions with significantly decreased errors. **(B)** Snapshots of IGV tracks for tRNA alignments. **(C)** The percentage of deletions and mismatches at each position of tRNA iso-decoders were extracted and summarized from the .mpileup file. * *p* < 0.05. Data are presented as mean ± SD. N=2. See also Figure S5 and S6 and Table S2.

tRNA-Tyr-GUA from human placenta and bovine liver is known to harbor two consecutive m^2^_2_G modifications at positions 26 and 27.^47,49^ While human TRMT1, the close relative of TRMT1L, has been demonstrated to catalyze the dimethylation at position G26, the enzyme responsible for catalyzing m^2^_2_G at position 27 has yet to be identified.^10^ The Nano-tRNAseq analysis detected the presence of m^2^_2_G26 and m^2^_2_G27 on tRNA-Tyr-GUA from WT cells, which appeared as deletions instead of mismatches at both positions. While the percentage of deletions was significantly decreased for both positions in TRMT1L KO cells, position G26 showed increased mismatch frequency, and position G27 showed an overall decrease in base calling errors (Figure 4A-C and Table S2). This profile indicates that two consecutive m^2^_2_G are detected as deletions at both positions and the loss of m^2^_2_G27, while maintaining m^2^_2_G26, is detected instead as increased mismatch frequency at position m^2^_2_G26 only. These findings suggest that TRMT1L is likely responsible for the deposition of m^2^_2_G27 on tRNA-Tyr-GUA. To validate this finding, we employed a primer extension assay capable of detecting RNA modifications with precise nucleotide resolution.^50–52^ This method can specifically identify m^2^_2_G, which impedes the elongation of the reverse transcriptase (RT) at the nucleotide carrying the modification,^53^ in this instance, position 27 (Figure 5A). As expected, we observed an RT stop at position G27 in WT cells, imposed by the m^2^_2_G modification reported at this position (Figure 5A). In contrast, this RT block was relieved in TRMT1L KO cells, which showed instead a stop at G26, a position known to harbor the m^2^_2_G catalyzed by the paralog TRMT1 (Figure 5A). Importantly, ectopic re-expression of TRMT1L, but not the catalytically inactive D373V mutant in the KO background restored the RT stop at position G27. No changes between WT and TRMT1L KO cells were detected for tRNAs Arg-UCG and Asn-GUU, both reported to contain m^2^_2_G26 catalyzed by TRMT1 (Figure S6B). A partial reduction, rather than complete loss, of m^2^_2_G on isolated tRNA-Tyr from TRMT1L KO cells was further validated via an immuno-Northern assay employing an anti-m^2^_2_G antibody. As expected, re-expression of TRMT1L WT, but not the catalytically inactive D373V mutant rescued the partial loss of m^2^_2_G detected by the antibody (Figure S6D). This analysis confirms the loss of m^2^_2_G at position 27 in TRMT1L KO cells, while position 26 retained the modification accounting for the remaining signal. Consistently, purified tRNA-Cys-GCA and tRNA-Arg-CCU were not detected by the anti-m^2^_2_G antibody, as previous reports have indicated the absence of m^2^_2_G modification in these tRNAs (Figure S6C).^18^

**Figure 5.**
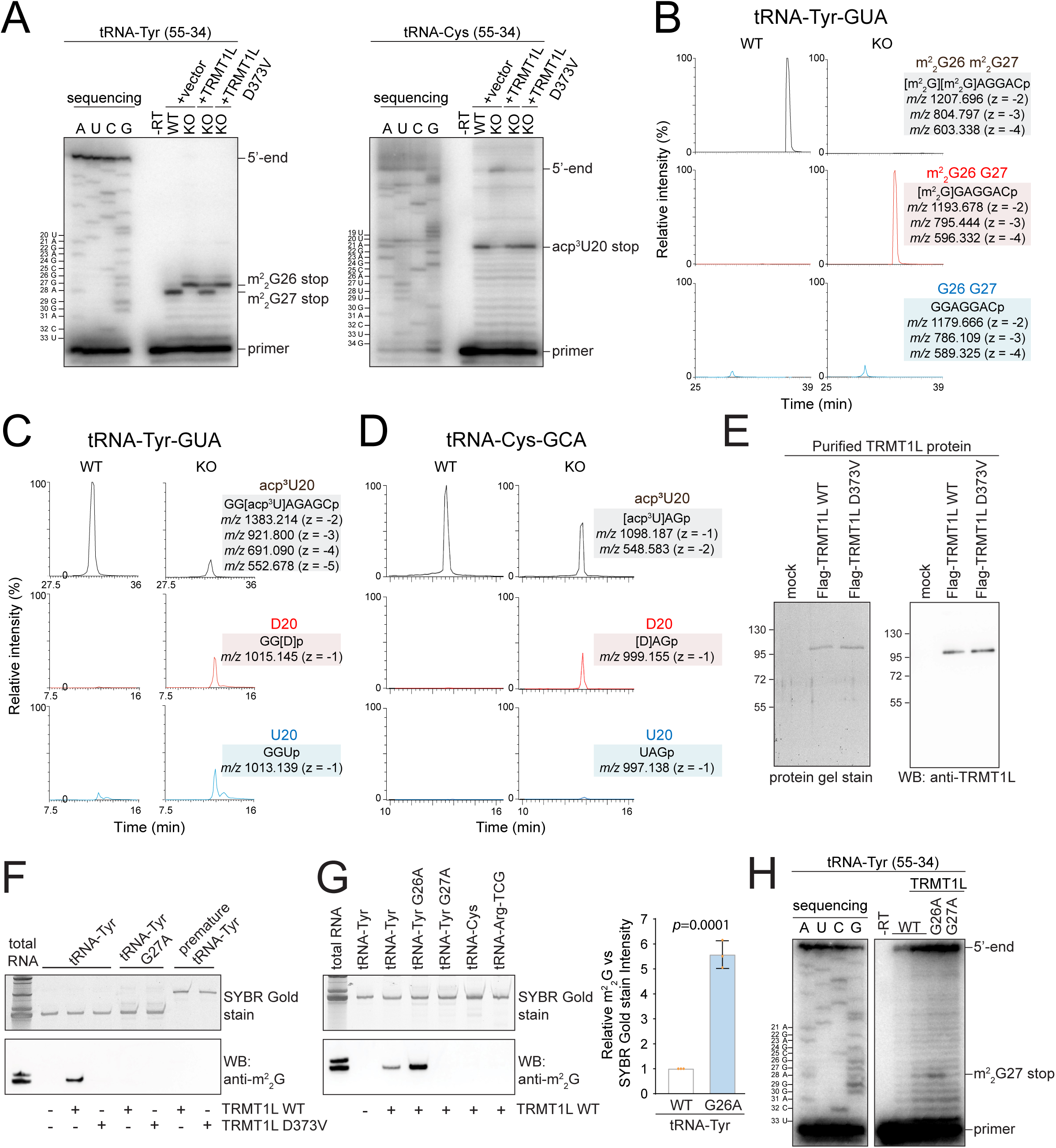
TRMT1L catalyzes the deposition of m^2^_2_G27 on tRNA-Tyr. **(A)** Primer extension analysis of tRNAs in WT, TRMT1L KO cells and in TRMT1L KO cells re-expressing TRMT1L WT or TRMT1L D373V mutant. **(B)** Extracted ion chromatograms of tRNA-Tyr-GUA RNase A digestion products. **(D)** Extracted ion chromatograms of tRNA-Cys-GCA RNase T1 digestion products. **(E)** Sypro Ruby protein gel stain and Western blot analysis of purified Flag-TRMT1L WT and TRMT1L D373V. **(F-G)** *In vitro* transcribed tRNAs were incubated with purified Flag-TRMT1L WT or D373V from (E) and analyzed by immuno-Northern blotting with m^2^_2_G antibody and densitometry quantification of the m^2^_2_G signals. Data are presented as mean ± SD. N=3. **(H)** The modifications on *in vitro* transcribed tRNA-Tyr WT, G26A and G27A after incubating with Flag-TRMT1L WT were analyzed by primer extension assay. Irrelevant lanes have been cropped. See also Figure S5 and S6.

The Nano-tRNAseq data also suggest loss of acp^3^U at positions 20 on tRNA-Tyr-GUA tRNA-Cys-GCA and tRNA-Ala-CGC/UCG in TRMT1L KO cells, which was detected as a decrease in mismatch errors in both cases (Figure 4A-C). tRNA-Cys-GCA was previously reported to lack m^2^_2_G at position 26 and our data did not identify any other potential modification showing a significant decrease in base calling errors in the TRMT1L KO cells for this tRNA. By primer extension assay, an RT block at position 20 on tRNA-Cys-GCA was observed in WT cells, confirming the presence of a modification likely to be acp^3^U20. In the TRMT1L KO cells, this RT block was substantially reduced (Figure 5A). Interestingly, re-expressed TRMT1L WT and the catalytically inactive mutant were both able to restore the acp^3^U20 RT block in the KO background. This finding indicates that TRMT1L does not catalyze, but is likely required to facilitate, the deposition of acp^3^U20 known to be catalyzed by DTWD1.^46^ The Nano-tRNAseq analysis further reveals that tRNA-Tyr-GUA appears to lack the D modification previously reported at position 17.^47,54^ In contrast, tRNAs Ala-CGC and Ala-UCG exhibit a less pronounced reduction of this reported modification at 16 in the TRMT1L KO cells (Figures 4A and 4C).^48^

To confirm the Nano-tRNAseq data, we isolated individual tRNAs Tyr-GUA and Cys-GCA and assessed their modification status using LC-MS/MS analysis of RNAse A (tRNA Tyr) or RNAse T1 (tRNA Cys)-generated fragments. For tRNA-Tyr-GUA, the fragment containing m^2^_2_G27 and m^2^_2_G26 observed in the WT cells was absent in the TRMT1L KO which showed instead a fragment containing only one m^2^_2_G modification predicted by the mapping software to be at position 26 (Figure 5B). Based on our Nano-tRNAseq and primer extension results, the fragment from the TRMT1L KO cells correspond to a loss of m^2^_2_G27 while m^2^_2_G26 is maintained. In the TRMT1L KO cells, the level of acp^3^U was drastically reduced on tRNA-Tyr-GUA, and to a lesser extent on tRNA-Cys-GCA. Instead, the loss or reduction in acp^3^U20 was accompanied by the modified bases converted to either D20 or U20 (Figure 5C-D). This is consistent with previous studies showing that acp^3^U formation inhibits D20 formation on tRNA-Tyr-GUA and Cys-GCA.^46^ Collision-induced dissociation spectrum was performed to confirm the modification identified on each fragment (not shown). Additionally, nucleoside analysis of purified tRNA-Tyr-GUA further confirms a reduction of m^2^_2_G and acp^3^U in the TRMT1L KO cells (Figure S6E and Table S3). The absence of D17 on tRNA-Tyr-GUA in TRMT1L KO cells requires further investigation, as we could not confidently confirm the presence of this modified position in the experimental condition tested. The acp^3^U modification on tRNAs Tyr-GUA and Cys-GCA is facilitated by the uridine aminocarboxypropyltransferase DTWD1, while DTWD2 is responsible for catalyzing acp^3^U deposition in tRNAs Asn-GUU, Ile-AAU, and Ile-UAU.^46^ Our Nano-tRNAseq did not reveal any decrease in the abundance of acp^3^U on tRNAs Asn-GUU, Ile-AAU, and Ile-UAU in TRMT1L knockout cells (Figure 4A), indicating that the depletion of TRMT1L specifically impacts the targets of DTWD1. In contrast, the dihydrouridine synthase DUS1L was recently shown to be responsible for introducing the D modification at positions 16 and 17 localized in the characteristic tRNA “D arm”.^54^

Finally, to demonstrate tRNA methyltransferase activity for TRMT1L, we conducted an *in vitro* reconstitution assay utilizing purified TRMT1L proteins, *S*-adenosyl-methionine, and *in vitro* transcribed tRNAs as substrates (Figure 5E-H). The assay revealed that purified TRMT1L WT, but not the catalytically inactive D373V mutant, catalyzes the formation of m^2^_2_G on tRNA-Tyr-GUA. This modification was detected via an immuno-Northern assay employing an anti-m^2^_2_G antibody and confirmed by primer extension assay (Figure 5F and H). Consistent with our eCLIP-seq data indicating a preference for intron-less tRNA-Tyr, TRMT1L deposited m^2^_2_G exclusively on mature tRNA-Tyr and not on its intron-containing precursor. No m^2^_2_G catalytic activity was observed for tRNA-Cys-GCA, which lacks this modification, nor for tRNA-Arg-TCG, which contains m^2^_2_G presumably catalyzed by the paralog TRMT1,^18^ thereby validating the specificity of our assay. To further ascertain specificity for G27 on tRNA-Tyr, we performed the assay with tRNA-Tyr substrates where G26 or G27 were substituted with A. TRMT1L did not exhibit any m^2^_2_G catalytic activity on the tRNA-Tyr G27A variant, whereas the tRNA-Tyr G26A variant showed about five times more m^2^_2_G levels compared to the WT tRNA-Tyr (Figure 5G). The reason for this is unclear but the G26A mutant could structurally expose G27 for m^2^_2_G catalysis byTRMT1L. These results were corroborated by primer extension assays (Figure 5H).

### TRMT1L-depleted cells have reduced levels of tRNA-Tyr-GUA and increased tRFs

To examine whether depletion of TRMT1L affects tRNA levels, we utilized the data from the Nano-tRNAseq analysis, a method shown to accurately quantify tRNA abundances.^55^ Normalized read counts of individual iso-decoders from both WT and TRMT1L KO samples showed a good correlation across biological replicates (Figure 6A-B). Compared to the WT cells, we observed a notable impact of TRMT1L depletion on the abundance of only one tRNA, namely tRNA-Tyr-GUA (Figure 6C). This nearly 2-fold reduction in tRNA-Tyr-GUA levels was validated through Northern blotting (Figure 6D-F). To determine whether the decrease in tRNA-Tyr-GUA level could be due to impaired structural stability due to its hypomethylation status, we analyzed RNA derived from WT and TRMT1L KO cells on native gel electrophoresis. The migration pattern of both tRNAs Tyr-GUA and Cys-GCA were unremarkably changed, indicating that the folding of these tRNAs was not dramatically altered or below the detection capacity of this method (Figure S7A). As tRNA modifications have been shown to regulate the binding and activity of aminoacyl-tRNA synthetases,^56–58^ we assessed the aminoacylation status of different tRNAs in WT vs. TRMT1L KO cell.^59,60^ Both Tyr-GUA and Cys-GCA, as well as a tRNA-Leu control from TRMT1L KO cells, appeared fully aminoacylated, demonstrating that the hypomethylated state of tRNAs Tyr-GUA and Cys-GCA does not affect amino acid charging (Figure S7B).

**Figure 6.**
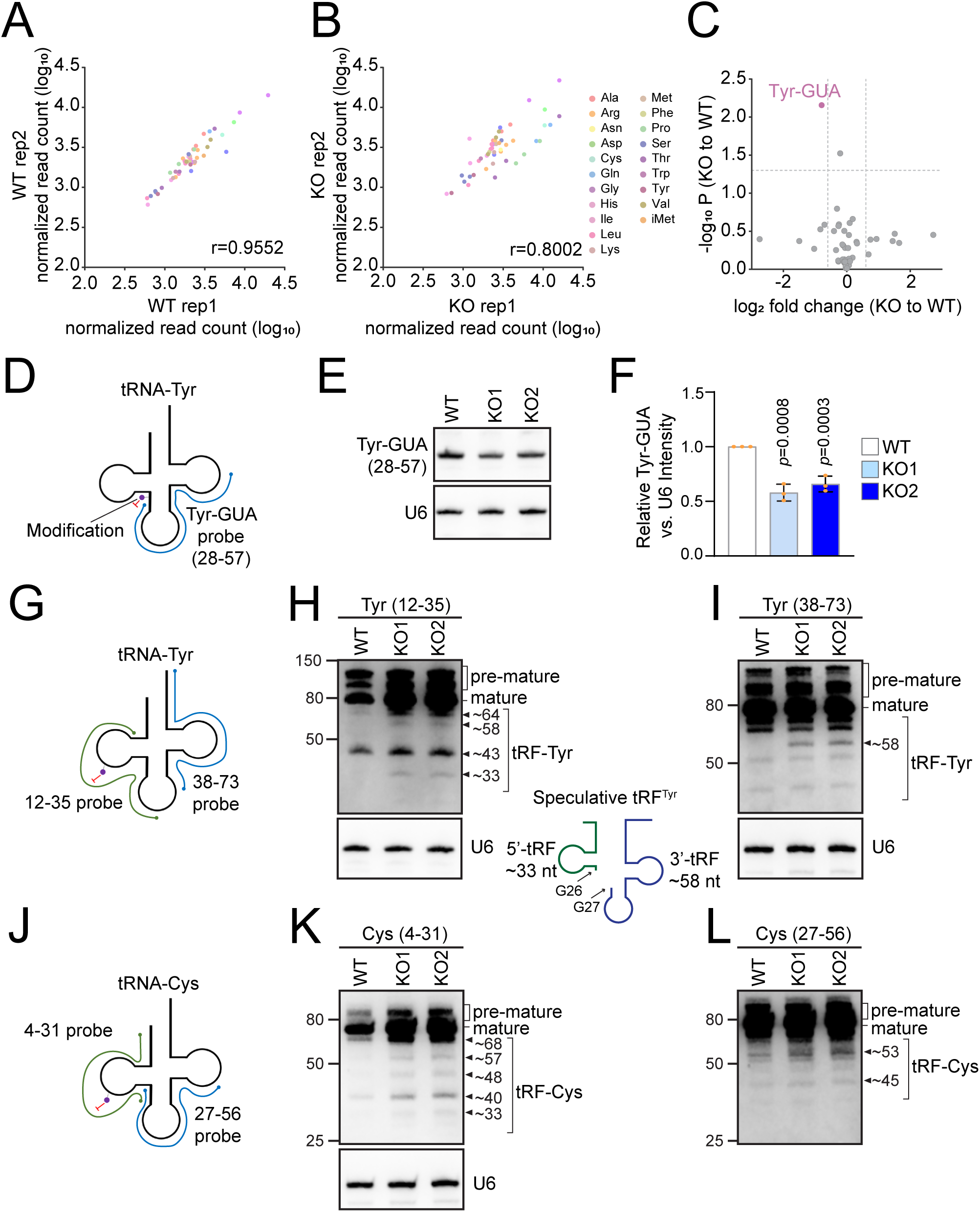
TRMT1L-depleted cells show a reduced abundance in tRNA-Tyr. **(A)** Nano-tRNAseq scatter plots of tRNA abundances of WT and **(B)** TRMT1L KO cells across biological replicates. The correlation strength is indicated by the Pearson correlation coefficient (r). **(C)** Volcano plot showing differential expression of tRNAs in TRMT1L KO relative to WT. The threshold of differential expression was set as *p* < 0.05 and absolute log_2_ fold change > 0.6. **(D, G, J)** Diagram showing the probes designed to cover different regions of tRNA-Tyr and tRNA-Cys. (**E**) Total RNAs from HCT116 WT and TRMT1L KO cells were analyzed by Northern blot using the indicated Tyr-GUA and U6 probes. **(F)** Densitometry quantification of Tyr-GUA signal normalized to U6 probe signal from (E) Data are presented as mean ± SD. N=3. **(H-I, K-L)** Total RNAs from HCT116 WT and TRMT1L KO cells were analyzed by Northern blot using the depicted tRNA-Tyr probes, tRNA-Cys probes and U6 probe. Approximate sizes of fragments detected in nucleotides are indicated on the side of each panel. Diagram showing the speculative fragments generated from a cut at position 26/27. The blots in (K-L) were done on the same membrane. See also Figure S7.

Interestingly, we observed that TRMT1L KO cells displayed an increased abundance of small tRNA-derived RNA fragments (tRFs), specifically for tRNA-Tyr-GUA and Cys-GCA (Figure 6G-L) and not for other tRNAs tested (Figure S7C). These fragments and the mature tRNAs appear at higher intensities using the Tyr (position 12-35) or Cys (position 4-31) probes in the TRMT1L KO compared to WT due to the PHA effect caused by the lack of m^2^_2_G27 and acp^3^U20. tRFs are generally products of precursors or mature tRNAs cleaved by different ribonucleases at specific positions.^61^ tRFs were observed in the TRMT1L KO cells, with probes specific for both the D and T-loop regions and ranged from approximately 35 to 55 nucleotides in length. Some of the most abundant fragments observed to be unique to TRMT1L KO cells could correspond to potential cleavage sites of the precursor tRNA-Tyr-GUA transcript at position 26/27 (fragments of 33 nt and 58 nt in length) (Figure 6H-I). Presumably, such fragments would originate from a spliced tRNA-Tyr-GUA precursor that still contains the 5’ leader and 3’ trailer sequences. While the precise nature of these observed tRFs needs to be confirmed by sequencing analysis, our data suggest that the absence of m^2^_2_G27 could render tRNA-Tyr-GUA more susceptible to cleavage at this position, possibly due to structural alterations, thereby contributing to its depletion in TRMT1L KO cells. Although we did not observe significant changes in tRNA-Cys-GCA levels, our findings indicate that depletion of TRMT1L also makes this tRNA more susceptible to cleavage potentially due to a reduction in acp^3^U20. It is also possible that TRMT1L may function as a chaperone, protecting tRNA from cleavage, which will need further investigation.

### TRMT1L is required for efficient mRNA translation and cell survival under oxidative and ER stress

The decrease in tRNA-Tyr-GUA levels and the elevated presence of tRFs observed in the TRMT1L KO cells prompted us to investigate whether the depletion of TRMT1L influenced translation and proliferation. Polysome profiling on sucrose density gradient reveals that cells depleted of TRMT1L slightly reduce mRNA translation efficiency, as observed by their lower polysome profiles compared to WT cells (Figure 7A). Because the TRMT1L KO cells were generated through puromycin selection, we used siRNA delivery to perform a puromycilation assay as a second means of measuring translation efficiency. Compared to control siRNA-treated cells, TRMT1L knockdown cells showed a noticeable reduction in global translation rate (Figure 7B), corroborating our polysome profiling observations. Surprisingly, given the decrease in mRNA translation observed, TRMT1L KO cells showed no significant proliferation defects compared to WT cells (Figure 7C).

**Figure 7.**
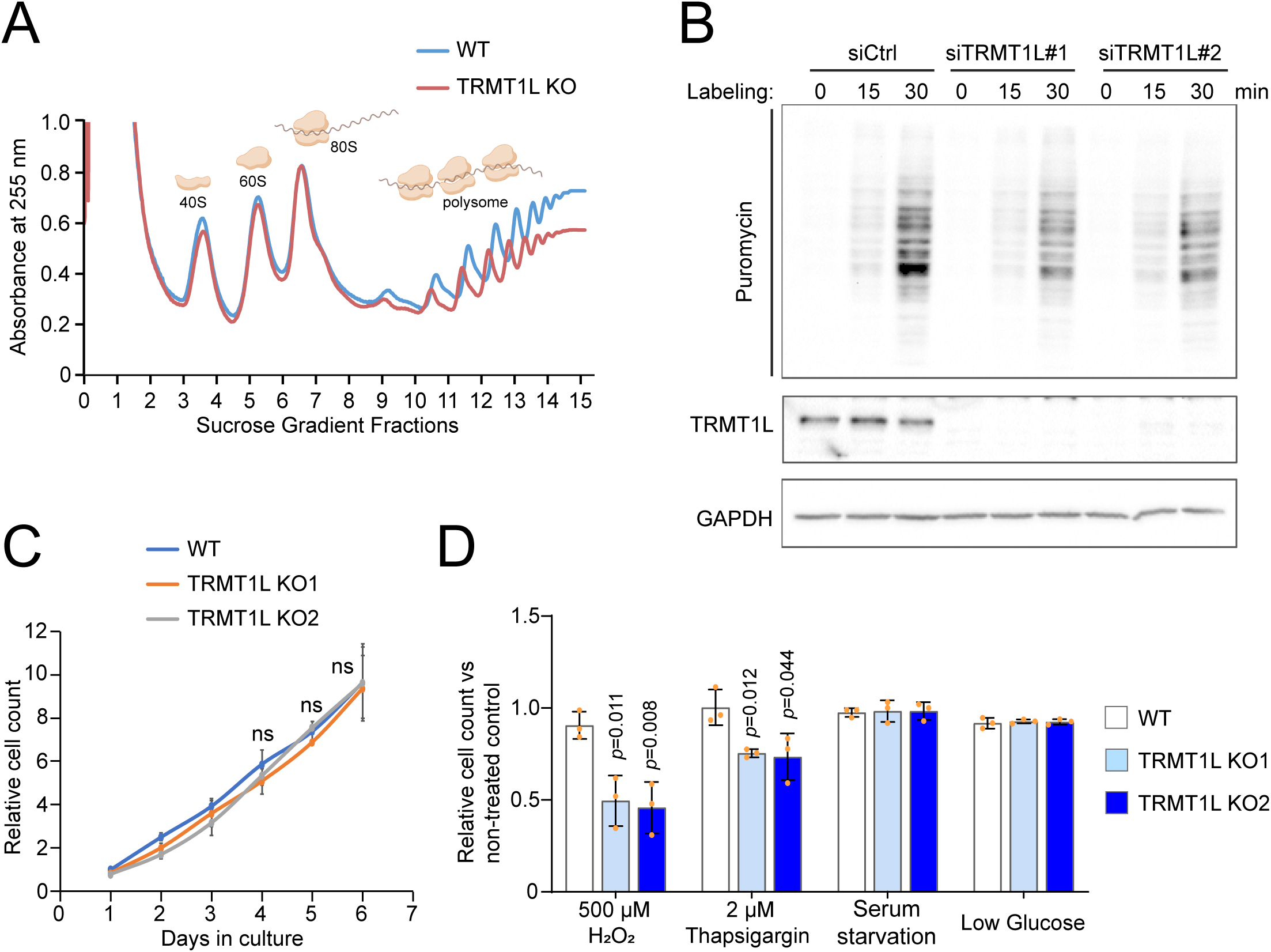
TRMT1L-depleted cells exhibit reduced global translation rate and increased sensitivity to oxidative and ER stresses. **(A)** WT and TRMT1L KO HCT116 cells were analyzed by polysome profiling. **(B)** HCT116 cells were transfected with siRNAs and treated with puromycin for the indicated time and analyzed by Western blot with the indicated antibodies. **(C)** WT and TRMT1L KO HCT116 cells were monitored for proliferation by cell counting. Relative cell counts were normalized to the cell counts on day 1. ns: *p* > 0.05. **(D)** WT and TRMT1L KO HCT116 cells were subjected to different stress conditions and live cells were counted after 48 h of treatment. Data are presented as mean ± SD. N=3.

The levels of m^2^_2_G have been shown to increase under oxidative stress conditions.^23,24,62,63^ Interestingly, human cells depleted of the paralog TRMT1 exhibit heightened sensitivity to oxidizing agents.^9^ To test whether TRMT1L also plays an important role in stress response, we subjected WT and TRMT1L KO cells to different stressors. Similarly to what was reported for TRMT1-depleted cells,^9^ TRMT1L KO cells also displayed hypersensitivity to oxidative stress (low dose of H_2_O_2_) as well as ER stress but not nutrient deprivation stresses (Figure 7D). These findings highlight the pivotal role of TRMT1L in maintaining the homeostasis of a specific subset of tRNAs by catalyzing m^2^_2_G and likely coordinating the deposition of acp^3^U20 and D17 modifications by DTWD1 and DUS1L enzymes, respectively.^46,54^ These functions are essential for efficient protein translation and the cellular response to oxidative and ER stress.

## DISCUSSION

The presence of two consecutive m^2^_2_G modifications, occurring at positions 26 and 27, is relatively rare and has been observed in only a few tRNAs, such as tRNA-Tyr purified from bovine liver and human placenta, as well as on *A. aeolicus* tRNA-Cys.^47,49,64^ While TRM1 has been identified as the enzyme responsible for catalyzing both m^2^_2_G26 and m^2^_2_G27 on *A. aeolicus* tRNA-Cys, human TRMT1 has been found to catalyze only m^2^_2_G26 on tRNA-Tyr,^10,64^ leaving the identity of the methyltransferase responsible for depositing m^2^_2_G at position 27 unresolved. In this study, we demonstrate that TRMT1L functions as the elusive methyltransferase catalyzing m^2^_2_G27 on human tRNA-Tyr-GUA. While our eCLIP-seq analysis revealed binding to a broader spectrum of tRNA and rRNA expansion segments, we found that TRMT1L exhibits catalytic activity only towards tRNA-Tyr-GUA. This observation suggests that TRMT1L may have additional catalytically independent functions depending on the RNA substrate.

Ensuring the proper folding of tRNAs into functional structures is vital for their role in translation. Recent investigations have unveiled additional roles for RNA-modifying enzymes beyond their catalytic functions, hinting at a dual role as tRNA chaperones.^65^ Recent studies in fission yeast have uncovered chaperone-like activities for TRM1 where both WT and catalytically inactive mutants have been shown to facilitate RNA strand annealing and dissociation, indicating a direct involvement in tRNA folding beyond the deposition of m^2^_2_G.^66^ Similarly, our study suggests a potential conservation of dual catalytic and chaperone functions for TRMT1L, whereby certain tRNAs may rely more heavily on one activity over the other. Our findings reveal distinct pools of substrates for TRMT1L: a group of modified substrates, primarily represented by tRNA-Tyr-GUA isoforms, and unmodified tRNAs (Cys, Arg, Asn, etc.) that may engage in chaperone-like interactions. Future work will be needed to experimentally test whether TRMT1L has a chaperone or tRNA folding function independent from catalytic activity.

Our quantification of tRNA abundance revealed that the levels of only tRNA-Tyr-GUA were affected by the depletion of TRMT1L, indicating that the presence of m^2^_2_G27 is critical for the stability of this particular tRNA. Numerous studies have shown the significant impact of m^2^_2_G presence within the hinge region on the conformational dynamics of tRNA molecules.^19–22^ The incorporation of dimethylguanosine at position 26 disrupts potential base-pairing interactions with cytidine, thereby playing a crucial role in stabilizing tRNA structure and preventing alternative folding conformations.^67^ Experimental evidence suggests that tRNAs devoid of m^2^_2_G26 exhibit notable structural alterations within their core architecture, characterized by a lack of base-stacking interactions and consequential loss of tertiary structural stability.^68^ In the case of tRNA-Tyr-GUA, the deposition of m^2^_2_G at position 26 would favor interaction with A44, while this modification at G27 would allow pairing with U43 in the stem region. The lack of m^2^_2_G27 is expected to compromise the structural integrity of tRNA-Tyr-GUA, potentially resulting in misfolding and subsequent degradation. While native gel analysis did not reveal significant conformational abnormalities in tRNA-Tyr from TRMT1L KO cells, further *in vivo* chemical probing is warranted for a finer assessment of the accessibility of the hinge region in tRNA-Tyr-GUA lacking m^2^_2_G27.

Our findings that TRMT1L is necessary for the deposition of acp^3^U and D modifications on tRNA-Tyr and Cys suggest its involvement in a potential tRNA modification circuit, wherein the introduction of modifications likely follows a specific order. Modification circuits on tRNAs have primarily been elucidated in bacteria and yeast, where modifications are typically installed in a sequential manner.^69,70^ For instance, pseudouridylation (Ψ) at position 55 of tRNA-Phe commonly occurs early during nascent tRNA maturation, facilitating the subsequent deposition of 2’-O-methylthymidine (Tm) at nucleotide 54, followed by the catalysis of m^1^A58.^70,71^ Similarly, in vertebrate tRNAs, studies using microinjected frog oocytes have revealed a sequential order of modifications on tRNA-Tyr. Alongside the early appearance of Ψ55, Tm54, and m^1^A58, traces of m^2^_2_G begin to emerge on the unprocessed tRNA-Tyr precursor. Upon leader and trailer removal, the intermediate precursor becomes fully m^2^_2_G modified, simultaneous with the appearance of acp^3^U and D, suggesting the potential requirement of TRMT1L for the deposition of acp^3^U and D modifications on tRNA-Tyr.^72,73^ Upon closer examination of our eCLIP-seq data, we discovered that TRMT1L exhibits binding affinity towards pre-tRNA-Tyr-GUA and Cys-GCA, which both contain 3’ trailer sequences. This observation suggests that TRMT1L binding is an early event in tRNA maturation. Interestingly, many of the TRMT1L eCLIP reads associated with tRNA-Tyr lacked the intron sequence but retained the 3’ trailer regions. Our *in vitro* methyltransferase assays also indicate that TRMT1L catalyzes the deposition of m^2^_2_G on tRNA-Tyr after intron removal as it had no catalytic activity on an intron-containing pre-tRNA-Tyr substrate. Interestingly, recent studies have shown that overexpression of DUS1L, which catalyzes the deposition of D16/17, suppressed 5’ and 3’ processing of tRNA-Tyr-GUA.^54^ Together, these findings suggest a precise timing of modification events regulating the processing of tRNA-Tyr.

Since TRMT1L does not exhibit catalytic activity towards tRNA-Cys-GCA and re-expression of both TRMT1L WT and a catalytically inactive mutant restores the loss of acp^3^U on tRNA-Cys in the TRMT1L KO background, our cumulative results indicate that it is not the m^2^_2_G modification itself, but rather the binding of TRMT1L to tRNA-Tyr or tRNA-Cys that is crucial for DTWD1 and DUS1L enzymes to catalyze the deposition of acp^3^U and D on these tRNAs. One possibility is that binding of TRMT1L to tRNA-Tyr or tRNA-Cys recruits DTWD1 and DUS1L through direct interactions. It is also conceivable that through chaperoning activity, TRMT1L could facilitate the folding of an open D-loop conformation required for subsequent recognition and binding of DTWD1 and DUS1L. Future experiments will be required to fully understand how TRMT1L mechanistically coordinates the deposition of acp^3^U and D on these tRNAs. A similar study by Zhang *et al*. converges with our findings, demonstrating TRMT1L’s role in catalyzing m^2^_2_G27 on tyrosine tRNAs. They also show TRMT1L’s involvement in maintaining acp^3^U modification on certain tRNAs, suggesting TRMT1L’s non-catalytic functions, possibly as a chaperone. Our collective studies strengthen the understanding of TRMT1L’s roles, particularly in m^2^_2_G27 modification and its potential involvement in an acp^3^U and D modification circuit.

It is becoming increasingly clear that different forms of stress can alter the modification pattern of tRNAs. Notably, studies in budding yeast exposed to oxidative stress have shown increased levels of 2’-O-methylcytidine (Cm), 5-methylcytidine (m^5^C), m^2^_2_G on tRNAs. Deletion of the enzymes catalyzing these modifications renders cells hypersensitive to oxidative stress, indicating the crucial role of such modifications in modulating tRNA function during stress.^62^ Notably, cells deficient in TRMT1 have demonstrated defects in redox homeostasis and heightened sensitivity to oxidizing agents.^9^ While we have not directly measured the redox state of TRMT1L-depleted cells, we have observed increased sensitivity to oxidative and ER stresses. Our findings suggest that the functions of TRMT1L are likely essential for orchestrating a translational response to cope with these stressors effectively. The significant reduction in tRNA-Tyr levels in TRMT1L KO cells suggests that translation defects may in part arise from inefficiencies in translating mRNAs enriched in this specific cognate codon. While the depletion of TRMT1L did not affect the levels of tRNA-Cys and tRNA-Ala, the loss of acp^3^U and D modifications on these tRNAs may impair their function, contributing to the observed decrease in mRNA translation. Considering its association with oxidative stress, it is conceivable that TRMT1L may become activated to methylate a broader range of tRNA substrates upon stress induction, although this hypothesis requires further investigation.

While the studies presented here focused on tRNAs, our CLIP-seq analysis also showed that TRMT1L binds to the 28S rRNA expansion segment ES7L, revealing a previously unexplored connection between the tRNA and ribosome biogenesis machineries. Expansion segments are extra sequences found in eukaryotic rRNA, which are believed to reflect the complex regulation of protein synthesis associated with organismal complexity.^74^ Previous studies suggested that expansion segments can serve as interaction platforms for ribosome-associated factors, binding with specific mRNAs, regulating translation fidelity, and controlling compartment-specific translation.^75–79^ ES7L, the largest eukaryotic expansion segment, contains a highly conserved core signature fold and long tentacle-like extensions that undergo rapid evolutionary changes.^80^ In yeast, the rRNA expansion segment ES7L was proposed to bind aminoacyl tRNA synthetases, potentially modulating translation efficiency.^76^ Additionally, under sublethal levels of oxidative stress, yeast 25S rRNA undergoes strand cleavage due to the binding of redox-active iron to ES7L, suggesting a potential role for this process in the adaptive oxidative stress response.^81^ As we did not observe posttranscriptional modifications of ES7L catalyzed by TRMT1L the interaction detected in the CLIP-seq analysis could be indirect or might reflect a distinct role of ES7L in regulating tRNA modifications in the nucleolus. Although previous studies have primarily focused on the roles of expansion segments in mature ribosomes, the extensions in ES7L could also assist in the recruitment of proteins to pre-ribosomal complexes or play a role in mediating complex formation between rRNA and nuclear proteins under stress conditions. Future investigation will explore the interactions of TRMT1L with pre-ribosomes in regulating tRNA and rRNA functions, providing insight into translation programs reliant on its activities for adaptive responses to stress.

### Limitations of the study

While our eCLIP-seq analysis, conducted in two distinct cell lines, has provided insights into the binding preferences of TRMT1L, with tRNA-Cys and Tyr emerging as predominant substrates, the observed spectrum of TRMT1L substrates may vary across different tissue types and environmental conditions, necessitating a more comprehensive investigation. Our findings suggest that TRMT1L’s activity and substrate selection could be influenced by tissue-specific factors and environmental cues, such as oxidative and ER stresses. Therefore, future studies should aim to broaden the scope of investigation by examining TRMT1L’s role in diverse tissue contexts and under various stress conditions. Furthermore, while our study highlights the significance of TRMT1L-mediated tRNA modification in regulating tRNA tyrosine levels and global mRNA translation rates, our approach did not fully capture the dynamic changes in translational landscape during cellular recovery from oxidative and ER stresses. To address this limitation, future research efforts will incorporate Ribo-seq and proteomics approaches to profile actively translated mRNA with high resolution. This will enable a more comprehensive understanding of the regulatory mechanisms orchestrated by TRMT1L in cells undergoing stress recovery.

## RESOURCE AVAILABILITY

### Lead Contact

Requests for further information and resources should be directed to and will be fulfilled by the lead contact, Catherine Denicourt.

### Materials Availability

Plasmids and cell lines generated in this study are available from the lead contact without restriction.

### Data and Code Availability

The datasets generated and/or analyzed during the current study are available in the GEO (Gene Expression Omnibus) data repository under accession numbers GSE266587 (eCLIP-seq) and GSE266588 (Nano-tRNAseq).

## ACKNOWLEDGMENTS

This work was supported by the National Institutes of Health (grant number R01-CA230746 to CD and R01-GM058843 to P.A.L.). Sseu-Pei Hwang received the CPRIT Fellowship in the Biomedical Informatics, Genomics and Translational Cancer Research Training Program (BIG-TCR) (CPRIT RP210045). The graphical abstract was created with BioRender.com.

## AUTHOR CONTRIBUTIONS

S-P.H. and H.L. performed most of the experiments with the help of K.B., X.H., C.H., H.M., A.S., and C.D. C.D., D.P., and P.A.L. guided the research. J.C. guided the bioinformatics analyses. C.D. and S-P.H. designed the experiments and wrote the paper with the input from all authors.

## STAR METHODS

### Experimental model and study participant details Cells

All the cells were obtained from the American Type Culture Collection (ATCC, Manassas, VA, USA) and cultured under 5% CO_2_ atmosphere at 37°C in high glucose Dulbecco’s modified Eagle’s medium (DMEM) supplement with 5% fetal bovine serum (FBS) and 100 U/ml penicillin, and 100 µg/ml streptomycin. All the cells were passaged by trypsin/EDTA digestion before reaching 80% confluency with a maximum of 2 months after thawing.

### Method details

#### Cell culture, RNA interference, proliferation assay and stress conditions

Synthetic short interfering RNA (siRNA) oligonucleotides (Sigma) were reverse transfected into the cells with Lipofectamine RNAi Max following the manufacturer’s instructions. For transfection, 20 nM of siRNA oligonucleotide was mixed with 6 µl of Lipofectamine RNAi Max per well (6-well plate). The suspension cells were then mixed with the siRNA transfection mixture and incubate under 5% CO_2_ atmosphere at 37°C. The sequences of the siRNAs utilized are listed in Table S4. For the proliferation assay, 10,000 cells were plated into wells of a 48-well plate, and cultured for 6 days. The cells in each well were counted every 24 h. For the cellular stress assays the following treatments were applied. For oxidative stress, cells were incubated with growth media supplemented with 0 or 500 µM of H_2_O_2_ for 30 min. The cells were washed twice with phosphate buffered saline (PBS) and incubated in regular growth media for 48h. For endoplasmic reticulum (ER) stress, the cells were treated with 0 or 2 µM Thapsigargin for 48h. For the low glucose condition, the cells were washed three times with PBS and incubated in glucose-free DMEM supplemented with 5% FBS and 2.5 mM glucose for 48h. The cells were washed three times with PBS and grown in serum-free DMEM for 48h for serum starvation. The cells were counted with trypan blue exclusion for all conditions at the 48h time point.

#### Immunoprecipitation and immunoblotting

For immunoprecipitation experiments, cells were lysed in ELB buffer (50 mM HEPES, pH 7.2, 250 mM NaCl, 2 mM EDTA, 0.5% NP-40) supplemented with protease inhibitors (aprotinin, leupeptin, AEBSF) on ice for 15 minutes. Lysates were then cleared by centrifugation at 22,000 × g for 10 minutes at 4°C, and the supernatant was incubated with 1 μg of normal rabbit IgG, anti-PELP1 antibody, or anti-TRMT1L antibody for 1 hour with rotation at 4°C. Protein G Sepharose beads were added for an additional 1 hour with rotation at 4°C. For immunoblotting, cells were lysed using RIPA buffer (25mM Tris-HCl pH 7.6, 150 mM NaCl, 1% NP-40, 1% Triton X-100, 1% sodium deoxycholate, and 0.1% SDS) and protease inhibitors on ice for 15 min. Lysates were cleared by centrifugation at 15,000 rpm for 10 minutes at 4°C. Protein concentrations were quantified using a BCA kit. Equal amounts of proteins were separated by SDS-PAGE and transferred to a 0.45 µm nitrocellulose membrane. The antibodies used for detection are listed in the reagent section.

#### Proteomic mass spectrometry analysis

HEK293T cells were lysed in ELB buffer (50 mM HEPES, pH 7.2, 250 mM NaCl, 2 mM EDTA, 0.5% NP-40) containing protease inhibitors (aprotinin, leupeptin, AEBSF) on ice for 15 min, and lysates were cleared by centrifugation at 22,000 × *g* for 10 min at 4°C. The supernatants were incubated with 2 μg of normal rabbit IgG or anti-PELP1 antibody for 1 h with rotation at 4°C. Protein G Sepharose beads were added for another 1 h with rotation at 4°C. Precipitated complexes were eluted in SDS–PAGE sample buffer and separated on a 10% SDS–PAGE. The gel was silver-stained and proteins present only in the anti-PELP1 lane were cut out, digested with trypsin, and analyzed by mass spectrometry on an Orbitrap Exploris 480 instrument (Whitehead Institute Quantitative Proteomic Core).

#### Generation of TRMT1L knockout (KO) cells

To establish CRISPR-KO cell lines, guide RNAs targeting TRMT1L were designed with CHOPCHOP.^82^ A non-targeting sgRNA control was used to establish cells labeled as WT: 5’-AACCTACGGGCTACGATACG-3’, KO1 targeting region: 5’-GTAGTTGTGAGAGTTTTGAG-3’, KO2 targeting region: 5’-TGTTACATCCGTAGTGACGC-3’. The oligo sequences, available in Table S4, were cloned into pLentiCRISPR v2 as described previously.^83,84^ To produce viral particles, the lentiviral vector containing the guide RNA was co-transfected to HEK-293T cells along with lentiviral packaging vectors, pMDL, pVSVG, and pREV in a ratio of 3:1:1:1.^85^ 24 h following transfection, the media was replaced, and the virus-containing media was collected at 48 and 72 h post transfection. The TRMT1L KO cells were generated by incubating HCT116 cells with the virus-containing media for 24 h. The infected cells were selected by puromycin. Gene deletion was assessed by DNA genotyping. Genomic DNA was extracted, and the TRMT1L targeted region was amplified with NEB Q5 polymerase and target-specific primers with Illumina adaptor overhangs. The PCR product was purified by gel extraction and sequenced on the Mi-Seq platform with 2×250 paired-end mode (Genewiz EZ Amplicon sequencing).

#### RNA extraction and Northern blot analysis

Total RNA was extracted using TRIzol reagent following manufacturer’s instructions. Total RNA (2-3 µg) was mixed with an equal volume of 2X urea-PAGE loading dye (2X: 95% formamide, 0.02% Bromophenol blue, 5 mM EDTA-NaOH (pH 8)) and denatured at 70°C for 5 min. Denatured RNAs were separated on a 12% denaturing (8 M urea) polyacrylamide gel followed by SYBR gold staining and visualization by transillumination at 302 nm UV. To analyze native tRNA structure, 2 µg of total RNA was combined with an equal volume of 2X native PAGE loading dye (2X: 20% glycerol, 0.012% bromophenol blue, 0.012% xylene cyanol, 10 mM MgCl2) and separated on 20% non-denaturing THEM (Tris/HEPES/EDTA/MgCl_2_) polyacrylamide gel at 4°C overnight as described by Woodson *et al.*^86^ For the pre-rRNA processing analysis, total RNA was separated on 0.8% formaldehyde denaturing agarose gel as described in Mansour *et al*.^87^ After electrophoretic separation, the RNA was transferred to an Amersham Hybond N+ nylon membrane and crosslinked to the membrane by 120 mJ/cm^2^ UV exposure. Northern blotting was performed as described previously.^88^ Briefly, membranes were pre-hybridized in hybridization buffer (0.6 M NaCl, 60 mM sodium citrate pH 7, 0.5% SDS, 0.1% BSA, 0.1% polyvinyl pyrrolidone, 0.1% Ficoll 400) at 50°C for 1 h and hybridized with 5 µM 5’-biotin-labeled probe in fresh hybridization buffer overnight. After hybridization and washes, probes were detected using streptavidin-alkaline phosphatase conjugate and Immun-Star AP substrate. The probes used for Northern blots are listed in Table S4. The membranes were stripped for other rounds of blotting by incubating the membrane in boiling stripping buffer (0.1X SSC, 0.1% SDS) twice. Densitometry analysis was performed using ImageJ.

#### Analysis of aminoacyl-tRNAs

The analysis of the aminoacylation status of tRNAs was performed according to protocols described in Janssen *et al.*^59^ and Zhang *et al.*^60^ RNAs extracted from both WT and TRMT1L KO cells were purified and reconstituted under acidic conditions to maintain the covalently bound amino acids. A subset of the RNA samples was subjected to deacylation under alkaline conditions to provide deacylated controls. Subsequently, the RNAs were separated via acidic polyacrylamide gel electrophoresis (PAGE), where aminoacylated tRNAs exhibit a slower migration rate than their deacylated counterparts. This difference in migration rate arises from the protonation of the α-amino group of the amino acid attached to the 3′ ends of the tRNA under acidic conditions. Briefly, total RNA was extracted with TRIzol and stored in acidic conditions to preserve aminoacyl-tRNAs (10 mM NaOAc (pH 5.0), 1 mM EDTA). As deacylated controls, a fraction of total RNA was subjected to deacylation under alkaline conditions in 50 µl of 100 mM Tris-HCl (pH 9.0), incubated at 37°C for 30 min and purified using the Monarch RNA Cleanup kit. 2.5 ug of RNA samples were combined with an equal volume of 2X acid loading dye (2X: 8 M urea, 0.1 M HOAc/NaOAc, pH 4.8, 0.05% bromophenol blue, 0.05% xylene cyanol), denatured at 95°C for 3 min and separated on a 12% acidic denaturing PAGE gel (8.3 M urea, 100 mM NaOAc (pH 5), 1 mM EDTA) in pre-chilled running buffer (100 mM sodium acetate (pH 5.0), 1 mM EDTA) at 16 W at 4°C for 24 h. The RNA was transferred to Amersham Hybond N+ nylon membrane and analyzed by Northern blotting as described in the section above.

#### Exogenous expression of wildtype, NLS mutant and NoLS mutant TRMT1L

To overexpress exogenous Myc-tagged TRMT1L, the TRMT1L coding sequence was cloned into pCMV-Myc. The nuclear localization signal (NLS) and nucleolar localization signal (NoLS) sequence of TRMT1L were predicted with cNLS Mapper^29^ and NoD tool,^27,28^ respectively. PCR mutagenesis was performed using the Q5 Site-Directed Mutagenesis Kit with primers listed in Table S4. Two lysine-to-alanine mutations (K621A and K622A) were introduced for the NLS mutant. For the NoLS mutant, R138A, R139A, H135A, and K136A mutations were made. Sequences of TRMT1L WT, NLS mutant, and NoLS mutant constructs were confirmed by Sanger sequencing (Genewiz). To re-express TRMT1L WT and TRMT1L D373V in the TRMT1L KO background for the rescue assays, the GFP coding sequence in pLenti-CMV-GFP-Hygro was replaced with TRMT1L WT or TRMT1L D373V cDNA. The mutation of TRMT1L D373V was achieved by PCR mutagenesis using the Q5 Site-Directed Mutagenesis Kit with primers listed in Table S4. Lentiviruses expressing GFP or TRMT1L were produced to infect TRMT1L KO cells. TRMT1L WT and TRMT1L D373V cDNA were also cloned into 3XFlag-CMV-7.1 for exogenous expression of Flag-tagged protein for *in vitro* methyltransferase assay.

#### Immunofluorescence staining

To localize endogenous TRMT1L by immunofluorescence, U2OS cells cultured on the cover glass and transfected with nontargeting scramble or TRMT1L siRNA for 72h were fixed with 4% formaldehyde and immunostained with anti-TRMT1L and anti-FBL (nucleolar marker) antibodies. Cells were treated with a DMSO vehicle or 500 nM CX-5461 to inhibit rRNA transcription for 1h before immunostaining. HCT116 cells transfected with Myc-tagged WT, NLS mutant, or NoLS mutant TRMT1L expressing plasmids on cover glasses were fixed with 4% formaldehyde and immunostained anti-Myc and anti-DDX21 (nucleolar marker) antibodies. The TRMT1L re-expressing HCT116 TRMT1L KO cells were fixed with 4% formaldehyde and immunostained anti-TRMT1L and anti-DDX21 antibodies. Primary antibodies were detected with Goat Anti-Mouse IgG H&L (Alexa Fluor 488) and Goat Anti-Rabbit IgG H&L (Alexa Fluor 594). Nuclei were stained with DAPI and visualized by fluorescence microscopy as previously described.^89^

#### Sucrose gradient density centrifugation for co-sedimentation assays

For the co-sedimentation assay, frozen HT1080 cells were incubated for 30 min on ice in lysis buffer (20 mM HEPES-NaOH pH 7.6, 150 mM NaCl, 0.5% Igepal CA-630, 1× protease inhibitor cocktail). The lysates were cleared by centrifugation at 18,400 × *g* for 20 min. The supernatant was loaded on a 10%-30% (wt/wt) sucrose gradient made in 10 mM HEPES-NaOH pH 7.6, 150 mM NaCl, 0.1 mM MgCl_2_, 1 mM DTT, and 0.01% Brij 35, and ultra-centrifuged at 36,000 RPM for 4 h at 4°C with a Beckman SW 41 Ti rotor. Each 1-ml fraction was thoroughly mixed, and RNA was isolated from 200 µl using TRI Reagent LS (Molecular Research Center, Inc), while the rest of each fraction was processed as described in Pestov *et al.,*^90^ to precipitate proteins for Western blot analysis. To assess defects in pre-ribosomal particle biogenesis, WT and TRMT1L KO cells were harvested and incubated in low salt buffer (10 mM HEPES-NaOH, pH 7.5, 10 mM NaCl, 2 mM MgCl2, 1 mM ethylene glycol tetraacetic acid) for 10 min and lysed by the addition of Igepal CA-630 and sodium deoxycholate to final concentrations of 0.3% and 0.2%, respectively as described in Pestov *et al.*^90^ Nuclei were sedimented by centrifugation at 1000 × *g* for 5 min and resuspended in sonication buffer (25 mM Tris-HCl, pH 7.5, 100 mM KCl, 1 mM NaF, 2 mM EDTA, 0.05% Igepal CA-630, 1 mM dithiothreitol [DTT]). Protease inhibitors (aprotinin, leupeptin, AEBSF) and RNase inhibitors were added, and nuclei were disrupted by sonication. The lysate was cleared by centrifugation at 15,000 × *g* for 15 min and was then separated on a 10–30% (wt/vol) sucrose gradient made in 25 mM Tris-HCl (pH 7.5), 100 mM KCl, 2 mM EDTA, and 1 mM DTT for 3 h at 160,000 × *g* using a Beckman SW 41 Ti rotor. Fractions were collected using a piston gradient fractionator (BioComp), and the absorbance at 255 nm was measured. Proteins in each fraction were extracted for Western blot analysis as described by Pestov *et al.*^90^

#### Polysome analysis

Polysome profiling was performed as described in Simsek *et al.*^91^ Cells were treated with 100 µg/ml cycloheximide for 5 min and lysed in polysome buffer (25 mM Tris–HCl pH 7.5, 150 mM NaCl, 15 mM MgCl_2_, 8% glycerol, 1% Triton X-100, 0.5% sodium deoxycholate, 1 mM DTT, 100 µg/ml cycloheximide, 100U/ml murine RNase inhibitor, 25 U/ml Turbo DNase, 1× protease inhibitor cocktail). The lysate was loaded on a 10-50% sucrose gradient and centrifuged at 40,000 rpm for 2.5h at 4°C. The fractions were collected by piston gradient fractionator (BioComp) and the absorbance at 255 nm was measured.

#### eCLIP-sequencing and biotin-labeling of immunoprecipitated RNA

The eCLIP-sequencing pipeline was adapted from procedures described by Van Nostrand et *al*.^31^ Cells were initially crosslinked with 400 mJ/cm^2^ UV on ice and lysed in CLIP buffer (50 mM Tris-HCl pH 7.4, 100 mM NaCl, 1% NP-40, 0.1% SDS, 0.5% sodium deoxycholate, 1× protease inhibitor cocktail) for 15 min. RNA fragmentation was accomplished through sonication, followed by treatment with RNase T1 and Turbo DNase at 37°C for 5 minutes. The nuclease-treated lysate was immunoprecipitated with protein G Dynabeads pre-bound with anti-TRMT1L antibody or rabbit IgG overnight. The co-precipitated complex was washed sequentially with high salt wash buffer (50 mM Tris-HCl pH 7.4, 1 M NaCl, 1% NP-40,0.1% SDS, 0.5% sodium deoxycholate, 1 mM EDTA) and wash buffer (20 mM Tris-HCl pH 7.4, 0.2% Tween-20). The RNA-ends were repaired on beads by treatment with Fast-AP and T4 PNK. For testing RNA binding activity of TRMT1L, a proportion of the dephosphorylated RNA was ligated to biotinylated cytidine (bis)phosphate (pCp-Biotin) by T4 RNA ligase. The biotin-labeled RNA samples were separated on an 8% SDS-PAGE gel, transferred to a nitrocellulose membrane, and bound RNA was detected using Streptavidin-IR800 on a LI-COR Odyssey DLx Imager. For eCLIP-seq, the rest of the dephosphorylated RNA was ligated to a 3’ RNA adaptor (/5Phos/rArGrArUrCrGrGrArArGrArGrCrArCrArCrGrUrC/3SpC3/) by T4 RNA ligase on beads. The co-precipitated complex and the samples for their respective paired size-matched input (SMInput) (two biological replicates for each sample) were resolved on an 8% SDS-PAGE gel and transferred to a nitrocellulose membrane for size selection. Samples ranging from the TRMT1L band to 75 kDa above on the membrane were excised, recovered, and prepared into paired-end libraries for high-throughput sequencing on an Illumina Hi-Seq 4000 platform (Novogene). Only samples from biological duplicates of TRMT1L immunoprecipitates and respective paired SMInput were processed for sequencing as the IgG control immunoprecipitated samples did not recover enough RNA to be amplified for library preparation. The raw fastq data were first trimmed to remove the adaptor sequence with CutAdapt.^92^ fastp was then used to remove and append the unique molecular identifier (UMI) to the read names.^93^ Reads shorter than 18 nt after trimming were discarded and the rest were aligned to the human genome (hg38) and human repetitive elements using the family-aware mapping approach developed by Van Nostrand *et al.*^32^ Reads mapping equally well to more than one family were discarded. Peak calling was conducted using the PEAKachu peak calling tool (https://github.com/tbischler/PEAKachu) within the pipeline, with parameters set as follows: --paired_end --max_insert_size 220. The deduplication was performed using UMI-tools^94^ with the parameter --read-length. Mapping profiles and statistics were extracted by Samtools.^95^ The numbers of reads identified as binding sites in the TRMT1L eCLIP samples were normalized to SMInput and significance and fold enrichment were calculated. To identify the crosslink site of TRMT1L on RNAs, the coverage of selected tRNA iso-decoders and rRNA genes were extracted from the reads aligned to the human genome (hg38) and analyzed with the mpileup function of Samtools. Reads in the TRMT1L eCLIP samples aligned to tRNA-Tyr-GTA-2-1 were further extracted and re-aligned to an intronless tRNA-Tyr-GTA-2-1 gene reference.

#### Puromycin labeling assay

HCT116 cells were transfected with 20 nM of non-targeting scramble or TRMT1L siRNAs for 72 hours and treated with 5 µg/ml of puromycin for 0, 15, and 30 min. Cells were harvested and processed for Western blotting analysis using an anti-puromycin antibody.

#### Purification of 28S rRNA

Total RNA was extracted from HCT116 WT or TRMT1L KO cells and resolved on a native 1% agarose gel. The separated RNAs were visualized under long-wave UV light, and the band corresponding to the 28S rRNA size was excised. RNA was then isolated from the gel slices using the Zymoclean Gel RNA Recovery Kit, following the manufacturer’s instructions. The purification of 28S rRNA was verified by resolving on 1% formaldehyde denaturing agarose gel, as described in Mansour *et al*.^87^

#### tRNA purification and single tRNA iso-decoder isolation

tRNAs were isolated from total RNA by ion exchange chromatography. The RNA samples were brought to a final concentration of 20 mM HEPES-KOH (pH 7.5) and loaded onto a Bio-Rad EconoFit Macro-Prep DEAE columns pre-equilibrated sequentially with the following buffers: 6 column volumes (CV) of low salt buffer (200 mM NaCl, 8 mM MgCl_2_, 20 mM HEPES-KOH pH 7.5), followed by 15 CV of high salt buffer (450 mM NaCl, 16 mM MgCl_2_, 20 mM HEPES-KOH pH 7.5), then 15 CV of low salt buffer, and finally, 15 CV of binding buffer (100 mM NaCl, 20 mM HEPES-KOH pH 7.5). After loading the RNA, the column was washed with 10 CV of binding buffer and eluted with a step gradient ranging from 200 mM NaCl, 8 mM MgCl_2_, 20 mM HEPES-KOH to 500 mM NaCl, 16 mM MgCl_2_, 20 mM HEPES-KOH. A final elution step was conducted with 1 M NaCl. After elution, the collected fractions were analyzed on urea-PAGE, and the fractions containing pure tRNAs were pooled and subsequently recovered by precipitation with 1/10 volume of 3M sodium acetate (pH 5.2), 20 ug/ml Glycogen and 1.5 volume of isopropanol.

Individual tRNAs were isolated from the purified tRNA using specific biotin-labeled DNA probes listed in Table S4. tRNAs were incubated with specific isoacceptor probes in binding buffer (0.5 M NaCl, 50 mM Tris-HCl (pH 7.5)) for 2 minutes at 98°C, followed by annealing at 80°C for 30 seconds and gradual cooling (−0.1°C/s) to 25°C. The annealed tRNAs were then captured on streptavidin magnetic beads, washed four times in binding buffer and eluted in RNase-free water. The remaining trace amounts of biotin-labeled DNA probes were removed by treating the eluate with Exonuclease I for 45 minutes at room temperature. This was followed by purification on the NEB RNA cleanup columns. The isolated tRNAs were analyzed for purity by Northern blot before further mass spectrometry analysis of modifications.

#### Mass spectrometry analysis of RNA modifications

For the oligonucleotide analysis, purified tRNA iso-decoders were digested with RNase T1 (50 U/ug) at 37°C for 1.5 hrs RNase A (0.01 U/ug) at 37°C for 1 hr. Analysis was done as previously described.^96^ The Synapt G2-S (Waters) coupled to a Dionex Ultimate 3000 RSLNano UHPLC (Thermo Scientific) was equipped with an Acquity UPLC BEH C18 column (1.7 μm, 0.3 × 150 mm, Waters). RNA digestion products were eluted at a 5 μl/min flow rate and 60°C column temperature with mobile phase A (8mM TEA and 200mM HFIP, pH = 7.8 in water) and mobile phase B (8mM TEA and 200mM HFIP in 50% methanol) with a linear gradient (3–55% of B over 70 min). Data was collected in sensitivity mode. MS scans were collected over 0.5 s in the m/z range of 545–2000, and MS/MS scans over 1 s period in the m/z range 250–2000. Three of the most abundant MS events were selected for MS/MS. A collision energy ramp (20–23 V at m/z 545; 51– 57 V at m/z 2000) was applied in a precursor m/z dependent manner, and the accumulated TIC threshold was set to 3.5 × 10^5^ counts. Oligonucleotide digestion products MS and MS/MS m/z were determined using the Mongo Oligo Mass Calculator v2.06 (http://rna.rega.kuleuven.be/masspec/mongo.htm).^97^ Data was collected and relative quantification was performed with MassLynx v4.2. Peak intensities are normalized to the sum of modified and unmodified digestion products.

For the nucleoside analysis, 28S rRNA or tRNA iso-decoders were digested to nucleosides using a previously described method.^98^ The RNA was denatured by heating at 95°C for 3 mins with an immediate cooling. The denatured RNA was incubated in 0.01 M ammonium acetate for 2 h with nuclease P1 (0.1 U/μg RNA) (Sigma-Aldrich). The RNA was then digested for 2 hr at 37 °C in 1/10 volume of 1 M ammonium bicarbonate with snake venom phosphodiesterase (1.2×10−4 U/μg RNA) and bacterial alkaline phosphatase (0.1U/ μg RNA) (Worthington Biochemical corporation). Analysis was performed on a TSQ Quantiva Triple Quadrupole (Thermo Fisher Scientific) coupled to a Vanquish Flex Quaternary (Thermo Fisher Scientific) with an Acquity UPLC HSS T3, 1.8 μm, 1.0 mm x 100 mm column (Waters). The nucleoside species were eluted at a 0.1 mL/min flow rate and 30 °C column temperature. Mobile phase A (5.3 mM ammonium acetate (pH = 4.5)) and mobile phase B (5.3 mM ammonium acetate, pH = 4.5, in 40% acetonitrile) were used with a previously described LC elution method.^99^ An SRM method was used for data collection to relatively quantify RNA modifications in samples using selected SRM transitions (Table S3). Analysis was done in positive polarity with an H-ESI source. The Q1 and Q3 resolution was 0.7, and a Dwell Time of 150 ms. For collision induced dissociation (CID), nitrogen gas was used at 1.5 mTorr achieving fragmentation. The sheath gas, auxiliary gas, and sweep gas of 35, 10, and 0 arbitrary units, respectively achieving; ion transfer tube temperature of 200°C; vaporizer temperature of 260 °C; and spray voltage of 3.5 kV was used for analysis. Data processing and relative quantification was done with Xcalibur 4.2 (Thermo Fisher Scientific).

#### Purification of Flag-tagged proteins

10 µg of p3xFLAG-CMV-TRMT1L or p3xFLAG-CMV-TRMT1L D373V was transfected to HEK293T cells. Two days after transfection, the cells were harvested and resuspended in lysis buffer (20 mM HEPES (pH 7.9), 2 mMgCl_2_, 0.2 mM EGTA, 10% glycerol, 1 × protease inhibitor cocktail) and sheered through a 26-gauge needle to lyse the cells. 0.3 M NaCl and 0.1% NP-40 was added after trituration. The mixture was incubated on ice and the lysates were cleared by centrifugation at 14,000 × *g* for 15 min at 4°C. The cleared lysate was mixed with 50 µl slurry of anti-DYKDDDDK Magnetic Agarose, and incubated at 4°C with rotation for 2 h. The agarose beads were washed three times with 0.3 M NaCl wash buffer (20 mM HEPES (pH 7.9), 2 mMgCl_2_, 0.2 mM EGTA, 10% glycerol, 0.1% NP-40, 0.3 M NaCl, 1 × protease inhibitor cocktail) followed by washing twice with 0.2 M NaCl wash buffer (20 mM HEPES (pH 7.9), 2 mMgCl_2_, 0.2 mM EGTA, 10% glycerol, 0.1% NP-40, 0.2 M NaCl, 1 × protease inhibitor cocktail), and once with 1X PBS. For elution of the purified protein, the beads were mixed with 50 µl of 1.5 µg/µl 3x DYKDDDDK Peptide in PBS and incubated for 10 min at room temperature with rotation at 1400 rpm. The elution was repeated once for maximum recovery. The purified proteins were aliquoted and stored at -20°C. The purified proteins were resolved on SDS-PAGE gel and stained by SYPRO Ruby Protein Gel Stain.

#### *In vitro* methyltransferase assay

To prepare the *in vitro* transcribed tRNA, templates were amplified from double-stranded DNA gBlocks (IDT) containing a T7 promoter sequence upstream of the tRNA sequence. This amplification was performed by PCR using the primers listed in Supplementary Table 4. The tRNA was transcribed *in vitro* from 50 ng of the PCR-amplified template using the MAXIscript™ T7 Transcription Kit. The transcription reaction mixture was incubated at 37°C for 1 hour, followed by TURBO DNase treatment at 37°C for 15 min, as the manufacturer’s instructions suggest. The transcribed RNA was purified using the NEB RNA Cleanup kit and resolved on a 12% denaturing (7 M urea) polyacrylamide gel, stained with SYBR Gold for visualization. Prior to the *in vitro* methyltransferase reaction, the transcribed RNA was refolded. For refolding, the RNA was first denatured in 5 mM Tris (pH 7.5) and 0.16 mM EDTA at 95°C for 2 min, then immediately cooled on ice for 2 min. The denatured RNA was refolded in a buffer containing 50 mM HEPES, 3 mM MgCl_2_, and 50 mM NaCl. For the methyltransferase reaction, 100 ng of refolded RNA was mixed with 25 nM of purified protein in a reaction buffer containing 50 mM Tris (pH 7.5), 0.1 mM EDTA, 1 mM DTT, and 0.5 mM *S*-adenosyl-methionine, and incubated at 30°C for 6 h. Post-incubation, the RNA was purified using the Monarch RNA Cleanup kit and resolved on a 12% denaturing (8 M urea) polyacrylamide gel, followed by SYBR Gold staining and visualization under 302 nm UV transillumination. The RNA was then transferred to an Amersham Hybond N+ nylon membrane and crosslinked by exposure to 120 mJ/cm² UV. The m^2^_2_G content in the crosslinked RNA was detected by immunoblotting with an m^2^_2_G antibody.

#### Primer extension assay

For preparing 5’-^32^P-labeled oligonucleotide, 20 pmol of oligonucleotide was incubated with 20 µCi [γ-^32^P]ATP and 10 U of T4 polynucleotide kinase in 1× T4 polynucleotide kinase reaction buffer at 37°C for 45 min. The reaction was heat-inactivated for 10 min at 68°C. Labeled primers were purified on a microspin G50 column (Cytiva 27-5330-01) per the manufacturer’s protocol. For the reverse transcription reaction, 3 µg of total RNA was mixed with 0.6 pmol of the 5’-^32^P-labeled oligonucleotides (Table S4) in 1X AMV Reverse Transcriptase buffer and incubated at 95°C for 3 min and slowly cooled to 42°C. 1.6 nmol of dNTP mix and 10 U of AMV reverse transcriptase were added to the reaction and incubated at 42°C for 1 h. The reaction was stopped by adding an equal volume of 2X urea-PAGE loading dye (2X: 95% formamide, 0.02% Bromophenol blue, 5 mM EDTA-NaOH (pH 8)). The corresponding sequencing lanes were prepared with *in vitro* transcribed tRNA as described in the “*in vitro* methyltransferase assay” section following the protocol developed by Nilsen *et al.*^100^ The reverse transcribed product was separated on a 15% urea-PAGE gel followed by fixation (10% methanol and 10 acetic acid). The gel was dried on a Whatman paper and exposed to a phosphorimager screen or x-ray films.

#### Nano-tRNAseq

The tRNA samples purified from total RNA were prepared into libraries following the Nano-tRNAseq protocol developed by Lucas *et al.*^36^ Initially, the tRNAs were deacylated in 100 mM Tris-HCl (pH 9.0) at 37°C for 30 minutes. After recovery using the NEB RNA Cleanup kit, the tRNAs were ligated to pre-annealed 5’ and 3’ splint adaptors (5’ RNA splint adapter: 5’-rCrCrUrArArGrArGrCrArArGrArArGrArArGrCrCrUrGrGrN, 3’ splint adapter: 5’-Phos-rGrGrCrUrUrCrUrUrCrUrUrGrCrUrCrUrUrArGrGrArArArArArArArArArAAAA-3’) with T4 RNA 2 Ligase followed by purification on AMPure RNAClean XP beads. The splint adaptor-ligated tRNAs were reverse transcribed using SuperScript IV Reverse Transcriptase at 60 °C for 1 hour followed by 85 °C for 5 minutes to linearize the tRNAs. The linearized tRNAs were cleaned up with AMPure RNAClean XP beads and then prepared into libraries following the standard protocol of the Direct RNA Sequencing Kit (SQK-RNA002) before being sequenced on a MinION flow cell (FLO-MIN-106). The bulk files were saved during sequencing for subsequent simulation runs using the alternative configuration described by Lucas *et al.*^36^ Fastq files generated by the simulation run were aligned to high confidence mature tRNA sequences retrieved from GtRNAdb ^101^ using the bwa alignment tool with parameters set as follows: -W 13 -k 6 -x ont2d -T20 ^102^. Sequence alignments in bam files were extracted by BEDTools.^103^ One tRNA gene was selected for each iso-decoder based on the read counts in this alignment to generate a reference file containing only one gene per tRNA iso-decoder. Differential expression of tRNAs was analyzed by aligning the fastq files to the one iso-decoder reference with the bwa alignment tool, using parameters set as follows: -W 13 -k 6 -x ont2d -T20, and extracting the alignment read counts by BEDTools. Mismatches, insertions and deletions at each position of tRNA iso-decoders were extracted from the sequence alignments in bam files using the mpileup function of Samtools ^95^. The percentage of deletions, mismatches and insertions of each position of tRNAs were summarized from the mpileup file. The sum of base calling errors for each tRNA iso-decoders position in TRMT1L depleted cells relative to WT was calculated by adding the rate of deletions, insertions and mismatches. An arbitrary weight of 1 for deletions and insertions and 0.6 for mismatches was assigned to identify tRNA modifications causing deletions. Sequence alignments were visualized using Integrative Genomics Viewer (IGV).^104^ Positions with a mismatch frequency greater than thresholds of 0.2 (Ala-CGC-2-1), 0.3 (Tyr-GTA-2-1), and 0.35 (Cys-GCA-3-1) are colored, whereas those lower than the set thresholds are shown in gray.

#### Quantification and statistical analysis

Data are shown as mean ± standard deviation. All data were generated from at least three biological replicates unless indicated. The eCLIP-seq and NanotRNA-seq experiments were performed for two biological replicates. Each dot on dot-bar plot represents an independent biological replicate. A two-tailed Student’s t-test determined the difference between the two groups. *p* value < 0.05 were considered to be statistically significant. All the Western, immuno-Northern and Northern blots were quantified using Image J.

## Supplemental Information

Document S1. Figures S1–S7 and Table S3.

Table S1. Excel file containing additional data too large to fit in a PDF. TRMT1L eCLIP sequencing peaks with IP vs SMInput fold-change >= 2 and adjusted *p* value < 0.05. Related to Figure 3, S3 and S4.

Table S2. Excel file containing additional data too large to fit in a PDF. Basecalling errors of each position of tRNA in Nano-tRNAseq. Related to Figure 4.

Table S4. Excel file containing additional data too large to fit in a PDF. Sequence of primers and probes used in this paper. Related to STAR Methods

---

**Figure S1.**
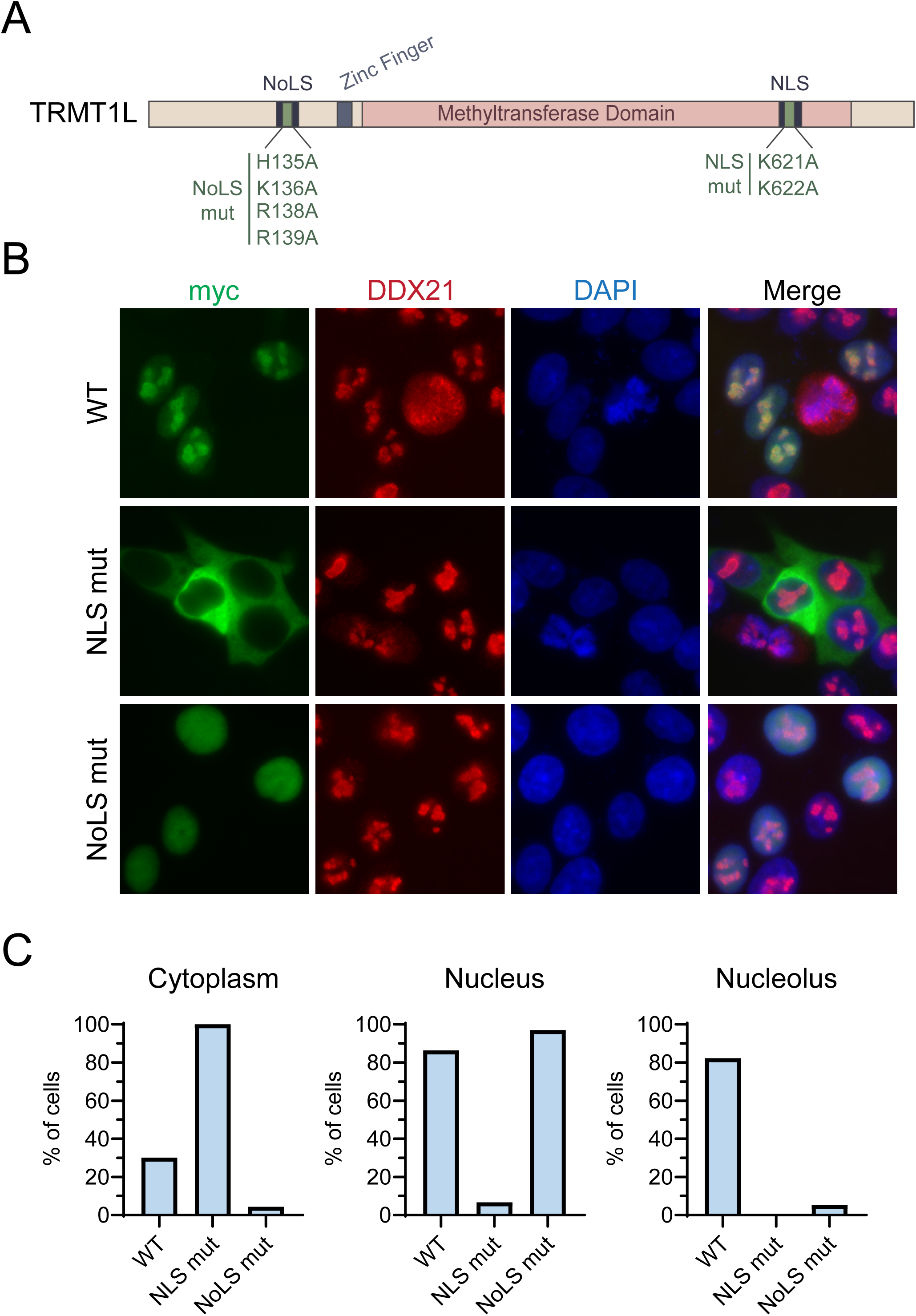
The subcellular localization of TRMT1L is dependent on its functional NLS and NoLS sequences. Related to Figure 1. **(A)** The schematic representation of TRMT1L protein with predicted sub-cellular localization signals and the indicated NoLs and NLS mutation sites. **(B)** HCT116 cells were transfected with Myc-tagged constructs of TRMT1L WT, NLS and NoLS mutants. The subcellular localization was detected by immunofluorescence with a Myc antibody. DDX21 was used as a nucleolar marker and nuclear DNA was stained with DAPI. **(C)** Quantification of the percentage of cells showing nuclear, nucleolar or cytosolic fluorescence signal for each construct from (B). 100 transfected cells were quantified for each construct.

**Figure S2.**
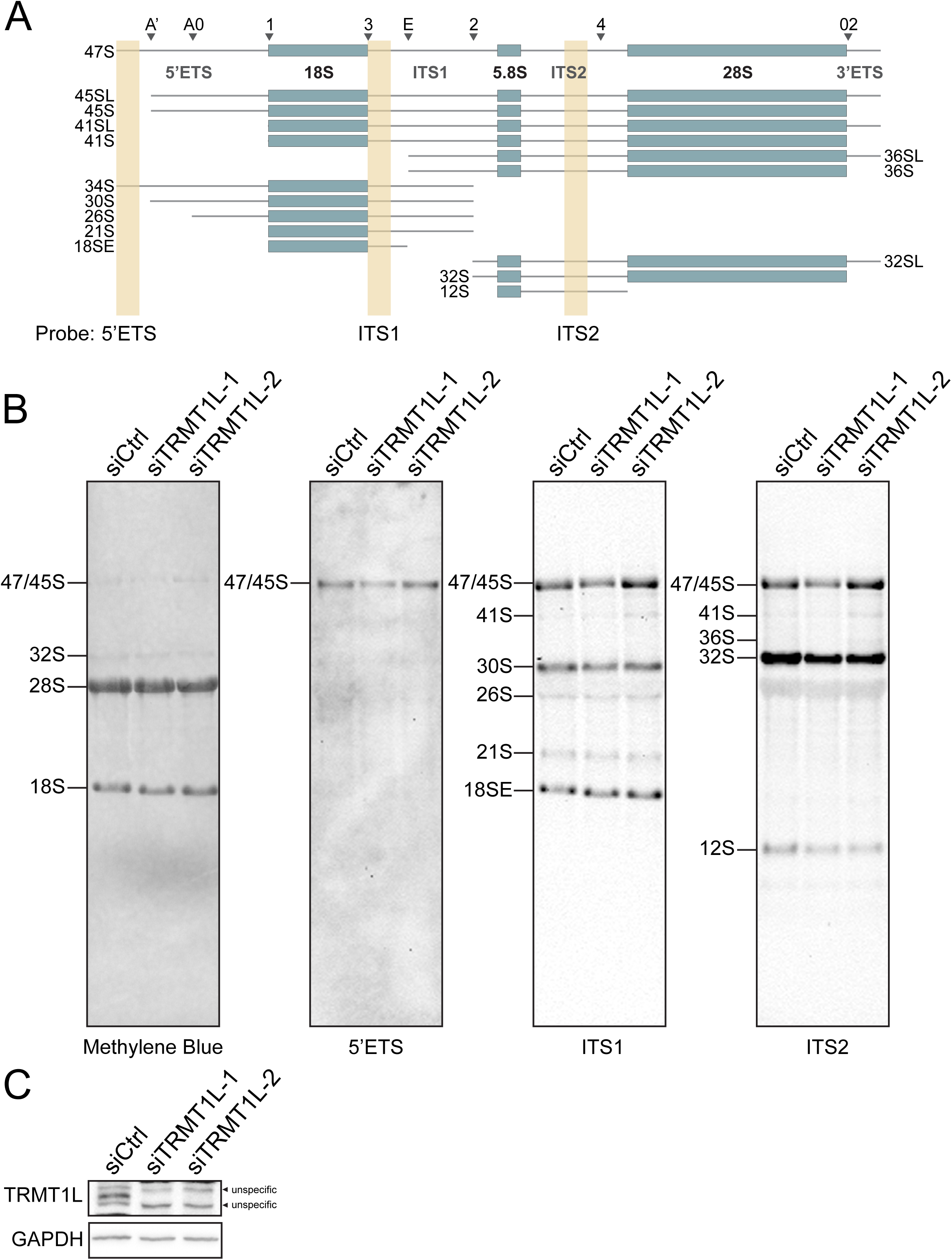
TRMT1L is not required for efficient pre-rRNA processing. Related to Figure 2. **(A)** Schematic overview of the human pre-rRNA processing pathway and the location of probes used for northern blotting are indicated by the cream-color-shaded area. **(B)** The total RNA of HCT116 cells transfected with non-targeting control (siCtrl) or TRMT1L siRNAs for 72 h were resolved on formaldehyde denaturing agarose gel and analyzed by Northern blot with probes targeting to 5’ETS, ITS1 and ITS2. **(C)** The TRMT1L expression in a fraction of cells from (B) was analyzed by Western blot with TRMT1L antibody.

**Figure S3.**
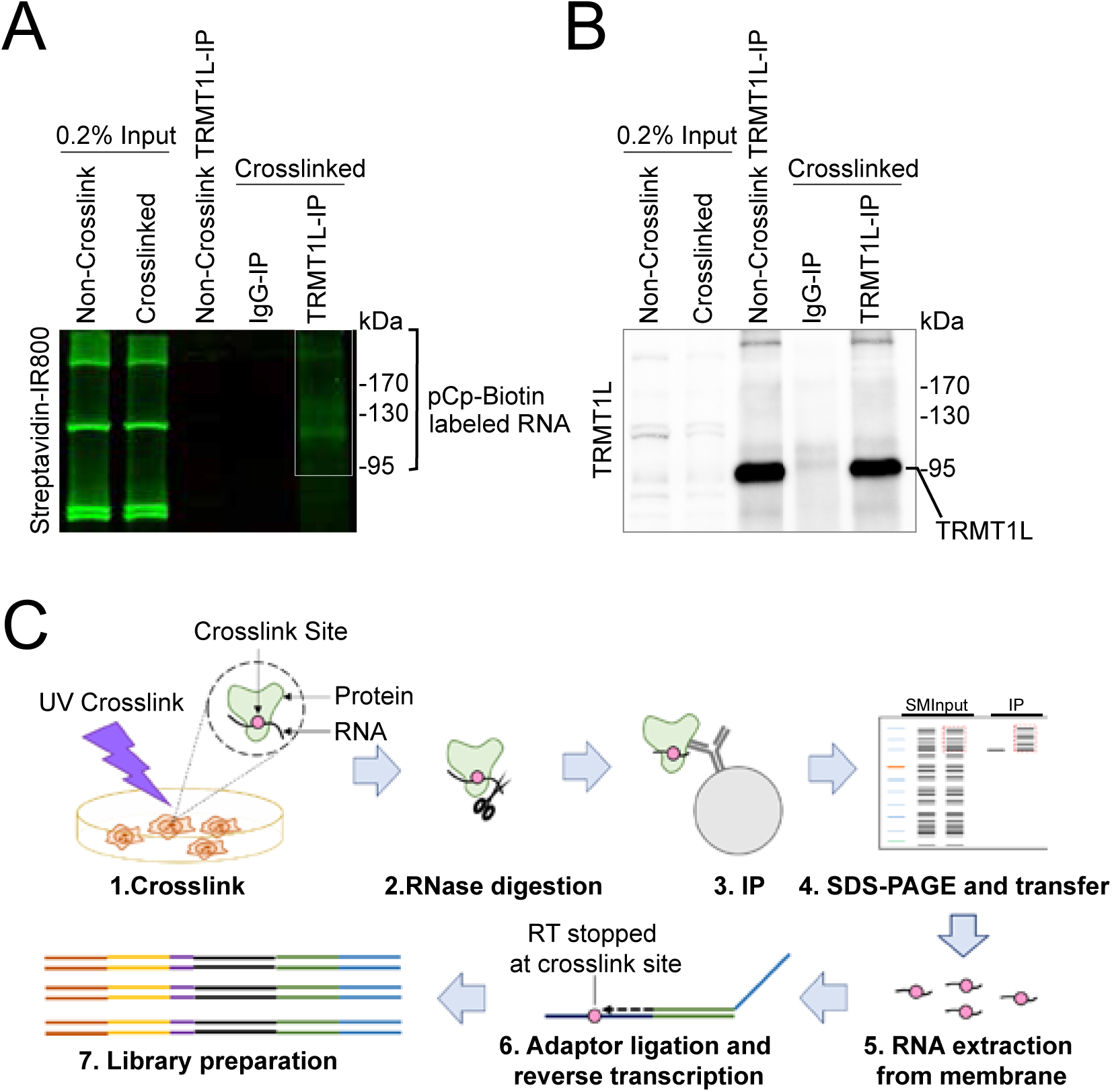
eCLIP-seq approach to identify TRMT1L RNA substrates. Related to Figure 3. **(A)** UV crosslinked and non-crosslinked HCT116 cells were lysed, digested with RNase, immunoprecipitated with TRMT1L or IgG control antibody, and 3’-end labeled with pCp-biotin for detection of co-precipitated RNA. After separation by SDS-PAGE and membrane transfer, the pCp-biotin labeled RNA was detected with Streptavidin-IR800 as indicated by the white box. **(B)** Immunoprecipitated TRMT1L was detected by immunoblotting with a specific antibody. **(C)** Schematic representation of the eCLIP-seq approach utilized to identify endogenous TRMT1L substrate.

**Figure S4.**
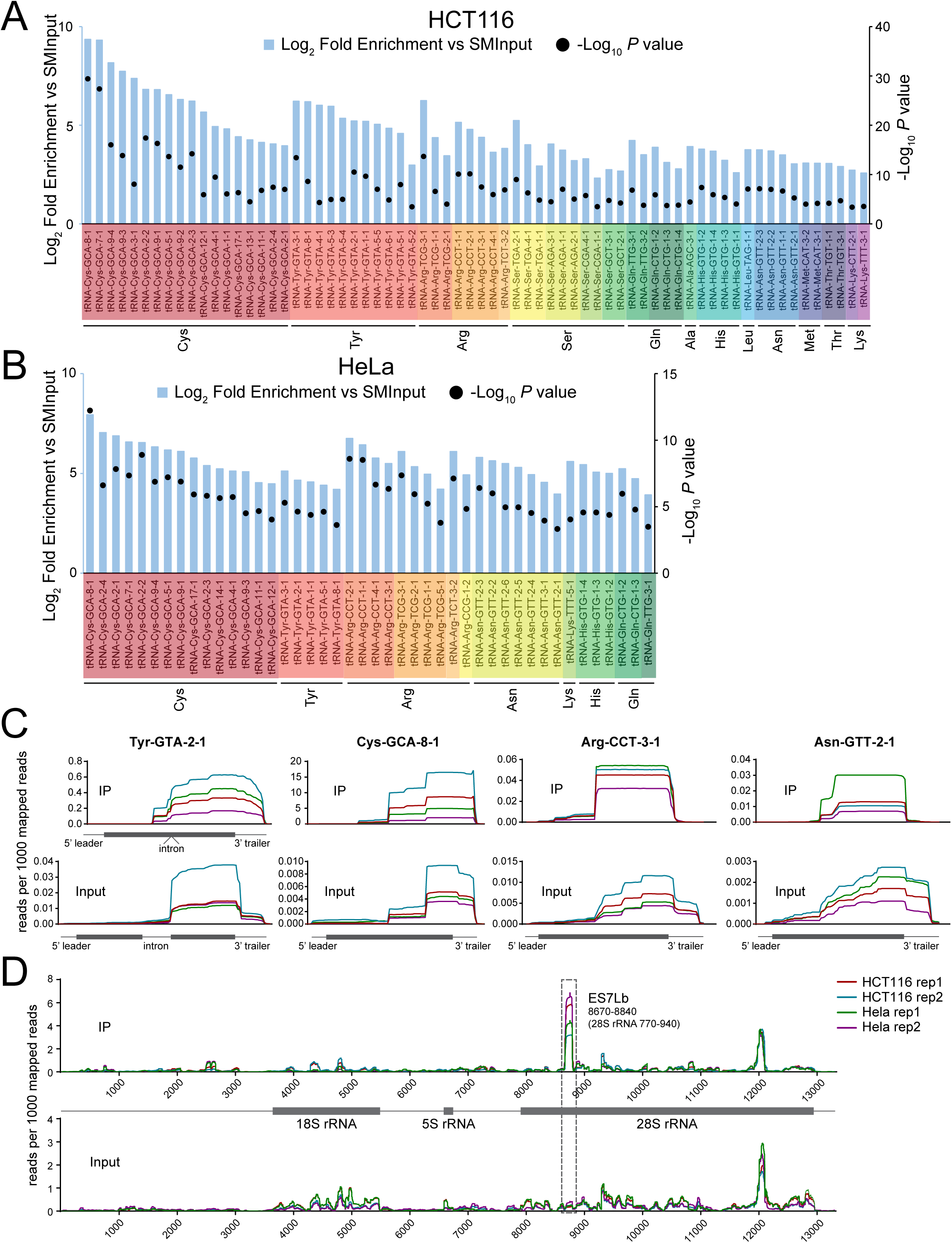
TRMT1L binds to tRNAs and rRNA. Related to Figure 3. **(A)** Complete list of tRNA iso-decoders identified by eCLIP-seq in HCT116 cells and **(B)** HeLa cells. Log2 fold enrichment of TRMT1L IP over size-matched input (SMInput) with cut-offs of adjusted significance p values < 0.05, fold change over size-matched input (SMInput) >= 2 and basemean >= 100. **(C)** The eCLIP-seq reads coverage of selected tRNA iso-decoders mapped on the whole genome. Reads in the IP samples aligned to tRNA-Tyr-GTA-2-1 were further extracted and re-aligned to an intronless reference to present the accurate tRNA enrichment. **(D)** eCLIP-seq reads coverage of human 47S rRNA (NR_146144.1 (RNA45SN2)). Dashed boxes mark the location of expansion segment ES7Lb. Coverage data was extracted from the whole genome alignment data using Samtools. Reads mapped to each gene were normalized to per 1000 mapped reads as indicated for IP and Input replicates. See also Table S1.

**Figure S5.**
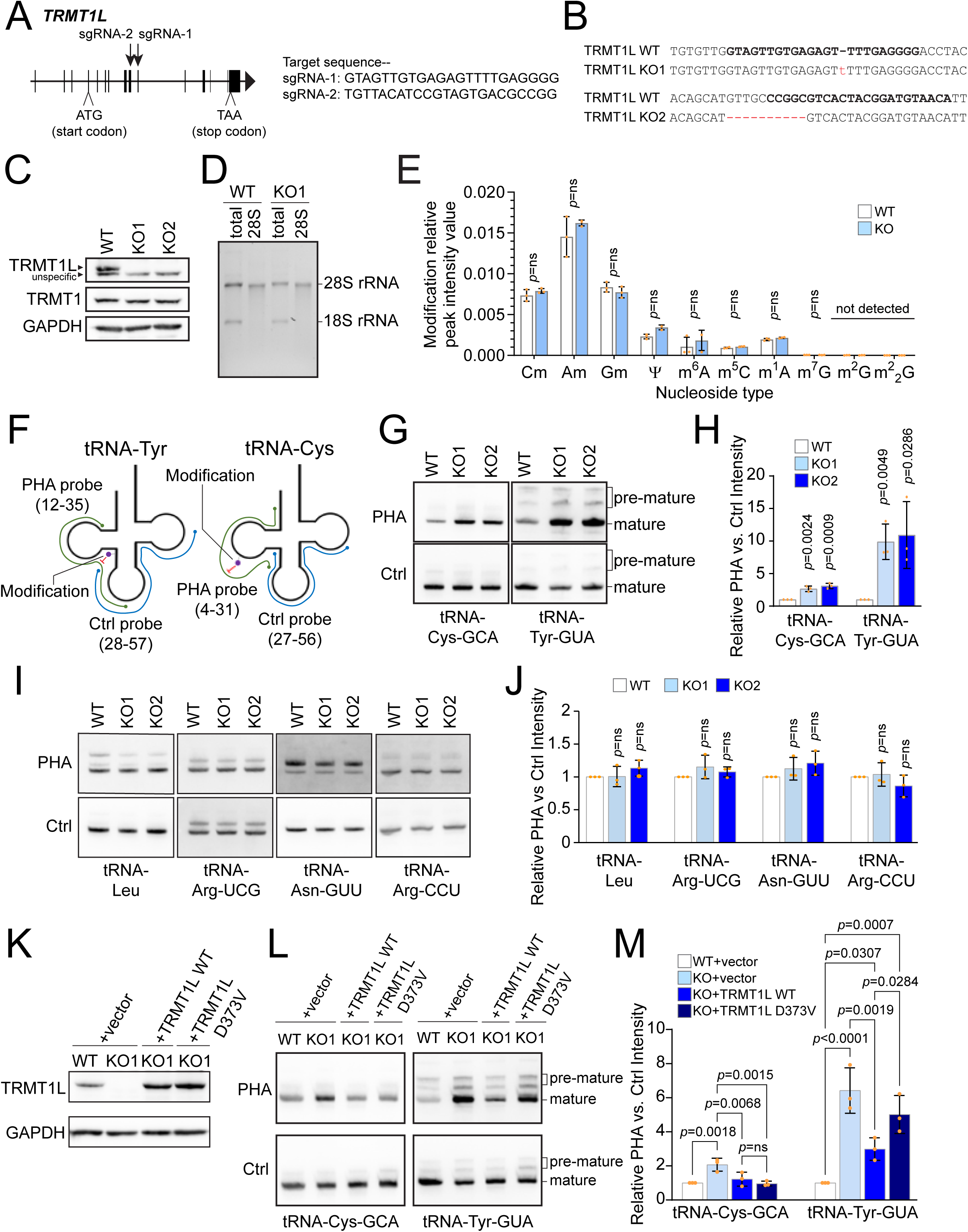
Depletion of TRMT1L is associated with loss of modifications on tRNAs Cys-GCA and Tyr-GUA. Related to Figure 4 and 5. **(A)** Diagram of the human TRMT1L genes and the designed guide RNAs (sgRNA) sequences and their targeting sites. **(B)** DNA genotypes of TRMT1L KO cell lines generated for each guide RNA by CRISPR Cas9. The targeting sequences are indicated in bold font, and the insertions and deletions in each are shown in red. **(C)** The TRMT1L and TRMT1 expression in HCT116 TRMT1L KO cells was analyzed by Western blot with TRMT1L and TRMT1 antibody as indicated. **(D)** The purification quality of isolated 28S rRNA from HCT116 WT or TRMT1L KO cells was verified by resolving on a 1% formaldehyde denaturing agarose gel. **(E)** Mass spectrometric nucleoside analyses of 28S rRNA modifications isolated from WT and TRMT1L KO cells. The relative peak intensity values of the indicated modifications of 28S rRNA from TRMT1L KO cells were normalized to WT. m^2^G and m^2^_2_G were not detected. **(F)** Positive hybridization in absence (PHA) of modification assay. The PHA probe covers the tRNAs 5’ half (D-loop region). A probe recognizing the 3’ half (T-loop region) was used as an internal control for normalization.**(G)** Total RNAs from HCT116 WT and TRMT1L KO cells were analyzed by Northern blot using the indicated probes as described in (F). **(H)** Densitometry quantification of PHA signal from (G) normalized to Control probe signal. **(I)** Total RNAs from HCT116 WT and TRMT1L KO cells were analyzed by Northern blot using the indicated probes. **(J)** Densitometry quantification of PHA probe signal from (I) normalized to Control probe signal. **(K)** Rescue re-expressions of TRMT1L WT or TRMT1L D373V in HCT116 TRMT1L KO cells were confirmed by immunoblotting with a TRMT1L antibody. **(L)** Total RNAs from HCT116 WT, TRMT1L KO and TRMT1L KO cells with exogenous re-expression of TRMT1L WT or TRMT1L D373V were analyzed by Northern blot using the indicated probes as described in (F). **(M)** Densitometry quantification of PHA signal from (L) normalized to Control probe signal. Data are presented as mean ± SD. N=3. Statistical significance between TRMT1L KO and WT PHA signal intensities was calculated using a 2-tailed independent student t-test.

**Figure S6.**
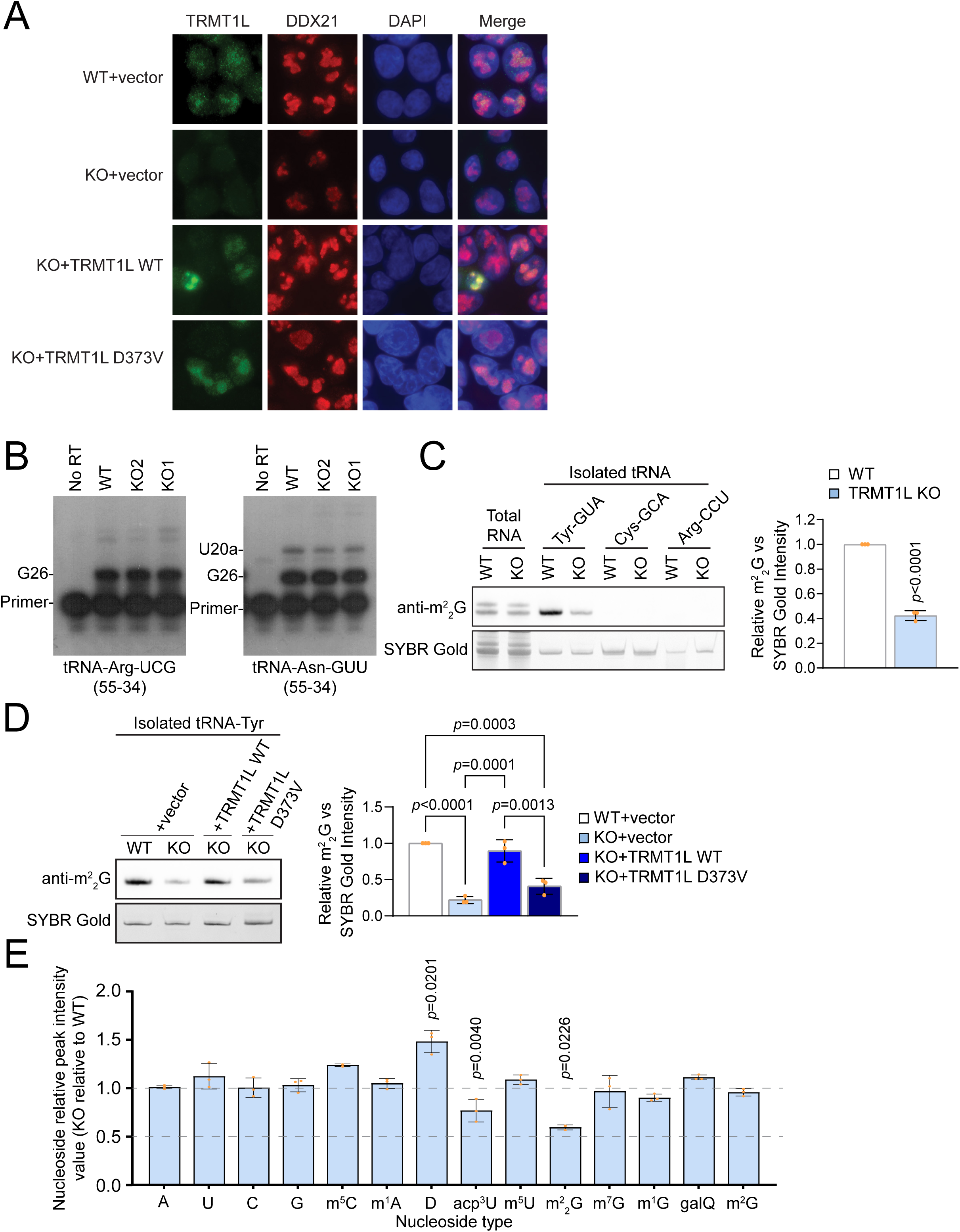
Depletion of TRMT1L is associated with loss of modifications on tRNAs Cys-GCA and Tyr-GUA. Related to Figure 4 and 5. **(A)** The immunofluorescence microscopy analysis showing nucleolar localization of TRMT1L WT or TRMT1L D373V re-expressed in HCT116 TRMT1L KO cells using a TRMT1L antibody. DDX21 was used as a nucleolar marker and nuclear DNA was stained with DAPI. **(B)** Primer extension assay showing no change of modification-induced reverse transcription (RT) block on tRNA-Arg-UCG and Asn-GUU in TRMT1L KO cells compared to WT. The position of RT primers for each tRNA is indicated in parentheses. **(C)** m^2^_2_G immuno-northern blot. The total RNA or isolated tRNA Tyr-GUA, Cys-GCA and Arg-CCU iso-decoders from the total RNA of HCT116 WT and TRMT1L KO cells were analyzed by immuno-Northern blotting with an m^2^_2_G antibody. Densitometry quantification of the m^2^_2_G signals normalized to SYBR gold signal for tRNA-Tyr isolated from WT and TRMT1L KO cells. **(D) I**solated tRNA Tyr-GUA iso-decoder from WT, TRMT1L KO, and TRMT1L KO cells re-expressing TRMT1L WT or D373V mutant were analyzed by immuno-Northern blotting with an m^2^_2_G antibody. Densitometry quantification of the m^2^_2_G signals was normalized to the tRNA SYBR gold signal. **(E)** Mass spectrometric nucleoside analyses of tRNA-Tyr modifications from WT and TRMT1L KO cells. The relative peak intensity values of the indicated nucleoside of isolated tRNA-Tyr from TRMT1L KO cells were normalized to WT. Data are presented as mean ± SD. N=3. See also Table S3.

**Figure S7.**
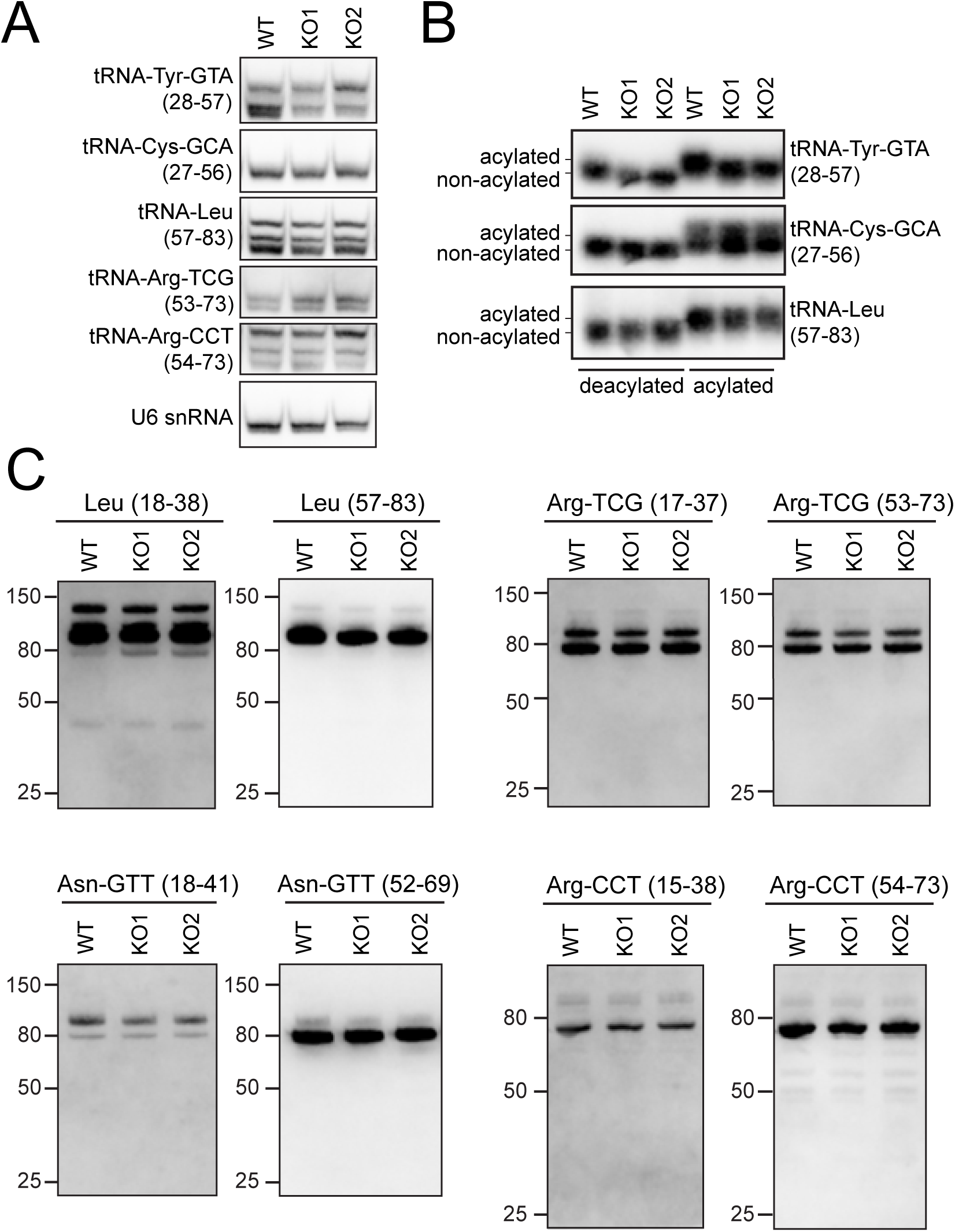
TRMT1L depletion does not affect the folding and amino acid charging of tRNAs and the level of tRFs derived from tRNA-Leu, tRNA-Asn-GUU, tRNA-Arg-UCG and tRNA-Arg-CCU. Related to Figure 6. **(A)** The total RNA from HCT116 WT and TRMT1L KO cells were separated on native PAGE and detected by Northern blot with the indicated tRNA probes. **(B)** The deacylated total RNA control and non-deacylated total RNA from WT and TRMT1L KO cells were resolved on 12% acid urea-PAGE gel and analyzed by Northern Blot using the indicated tRNA iso-decoder probes. **(C)** The total RNA from HCT116 WT and TRMT1L KO cells were analyzed by Northern blot with the indicated tRNA probes. For each tRNA, blots were consecutively hybridized with D-loop and T-loop specific probes. The blots are shown at a high exposure to detect the presence of tRFs.

**Table S3.** The selected SRM transitions of the RNA modifications used in nucleoside analysis. Related to Figure S6.

| **Compound** | **Precursor (m/z)** | **Product (m/z)** | **Collision Energy (V)** | **RF Lens (V)** |
| --- | --- | --- | --- | --- |
| **D** | 247 | 115 | 10 | 43 |
| **m2,2G** | 312 | 180 | 15 | 60 |
| **acp3U** | 346 | 214 | 10 | 58 |
| **C** | 244 | 112 | 12 | 41 |
| **U** | 245 | 113 | 10 | 30 |
| **m1A** | 282 | 150 | 20 | 68 |
| **m5C** | 258 | 126 | 14 | 46 |
| **G** | 284 | 152 | 14 | 46 |
| **m7G** | 298 | 166 | 16 | 50 |
| **m5U** | 259 | 127 | 10 | 30 |
| **galQtRNA** | 572 | 163 | 29 | 84 |
| **galQtRNA** | 572 | 295 | 29 | 84 |
| **m1G** | 298 | 166 | 16 | 50 |
| **A** | 268 | 136 | 17 | 63 |
| **m2G** | 298 | 166 | 16 | 50 |

